# Rat Deconvolution as Knowledge Miner for Immune Cell Trafficking from Toxicogenomics Databases

**DOI:** 10.1101/2023.06.20.545836

**Authors:** Katsuhisa Morita, Tadahaya Mizuno, Iori Azuma, Yutaka Suzuki, Hiroyuki Kusuhara

**Affiliations:** Department of Pharmaceutical Sciences, The University of Tokyo, Bunkyo, Tokyo, Japan; Graduate School of Frontier Sciences, The University of Tokyo, Chiba 277-8561, Japan

**Keywords:** deconvolution, toxicogenomics, immune cell, rat, transcriptome

## Abstract

Toxicogenomics databases are useful for understanding biological responses in individuals because they include a diverse spectrum of biological responses. Although these databases contain no information regarding immune cells in the liver, which are important in the progression of liver injury, deconvolution that estimates cell-type proportions from bulk transcriptome could extend immune information. However, deconvolution has been mainly applied to humans and mice and less often to rats, which are the main target of toxicogenomics databases. Here, we developed a deconvolution method for rats to retrieve information regarding immune cells from toxicogenomics databases. The rat-specific deconvolution showed high correlations for several types of immune cells between spleen and blood, and between liver treated with toxicants compared with those based on human and mouse data. Additionally, we found 4 clusters of compounds in Open TG-GATEs database based on estimated immune cell trafficking, which are different from those based on transcriptome data itself. The contributions of this work are three-fold. First, we obtained the gene expression profiles of 6 rat immune cells necessary for deconvolution. Second, we clarified the importance of species differences on deconvolution. Third, we retrieved immune cell trafficking from toxicogenomics databases. Accumulated and comparable immune cell profiles of massive data of immune cell trafficking in rats could deepen our understanding of enable us to clarify the relationship between the order and the contribution rate of immune cells, chemokines and cytokines, and pathologies. Ultimately, these findings will lead to the evaluation of organ responses in Adverse Outcome Pathway.

## INTRODUCTION

Immune cell trafficking is one of the most important factors in the progression of liver injury, such as drug-induced liver injury (DILI) (Yang and Tonnesseen, 2019; Jaeschke *et al*., 2012; Hossain and Kubes, 2019). While the importance of various immune cells, such as Kupffer cells and NKT cells, has been demonstrated individually, their contribution to the progression of liver injury and their relationship with each other have remained largely unknown (Liu *et al*., 2006; Laskin *et al*., 1995; Yang *et al*., 2019; Liu *et al*., 2004). One possible explanation for this situation is that it is difficult to compare accumulated findings due to the complexity of the experimental systems, such as the fact that liver injury models that have been constructed using various methods including compound administration, diet, and surgery, as well as different time points for evaluation (Cai *et al*., 2017; Gehring *et al*., 2006; Graubardt *et al*., 2017; Chen *et al*., 2017).

Large-scale toxicity databases of compounds (toxicogenomics databases), such as Open TG-GATEs and DrugMatrix, are useful for understanding biological responses in individuals (Igarashi *et al*., 2015), (https://ntp.niehs.nih.gov/data/drugmatrix). This is because compound administration is relatively easy to control and less confounded compared with other model preparation approaches such as diet and surgery, making it easier to compare findings. In addition, considering that compounds generally have multiple effects, including those unrecognized, it is also expected that a diverse spectrum of biological responses can be covered due to nonarbitrary perturbations by various compounds (Morita *et al*., 2020; Nemoto *et al*., 2021). Open TG-GATEs, for example, contain transcriptome data, pathological images, and blood biochemistry of tissues collected at various time points from rats treated with various compounds at various concentrations, while there is no information regarding immune cells in tissue in these databases.

Deconvolution is a data analysis method that can be applied to estimate the ratio of cells in a specimen from bulk transcriptome data (Abbas *et al*., 2009; Im and Kim, 2023). Bulk transcriptome is affected by distinctive sample conditions (e.g., gene knock out), individual or technical variations, and relative cell subset proportions. Based on this concept, it is possible to analyze the impact of proportion changes on bulk transcriptome, by focusing on gene subsets uniquely expressed in target cells. To elaborate this, cell proportions can be calculated through a regression model using bulk transcriptome of a specimen, transcriptome of target cells, and specific marker genes for the target cells. Applying this method to transcriptome data of a toxicogenomics database, could add information regarding immune cell trafficking as a new layer to the database and contribute to a deeper understanding of immune cell relationships in liver injury. However, deconvolution has been mainly applied to humans and mice, and there have been few applications to rats, which are the main target of large-scale toxicity databases (Z. Chen *et al*., 2018; Altboum *et al*., 2014; Wang *et al*., 2021; Petitprez *et al*., 2020). In particular, a gene expression dataset of various rat immune cells is currently absent and prior studies applying deconvolution to rats have used human immune cells as a substitute (Gil Del Alcazar *et al*., 2022; Wang *et al*., 2021).

Here, we developed a rat deconvolution method, examined species differences in the deconvolution method, and established a methodology to obtain immune cell information from valuable toxicogenomics databases (**Figure 1**).

**Figure 1.**
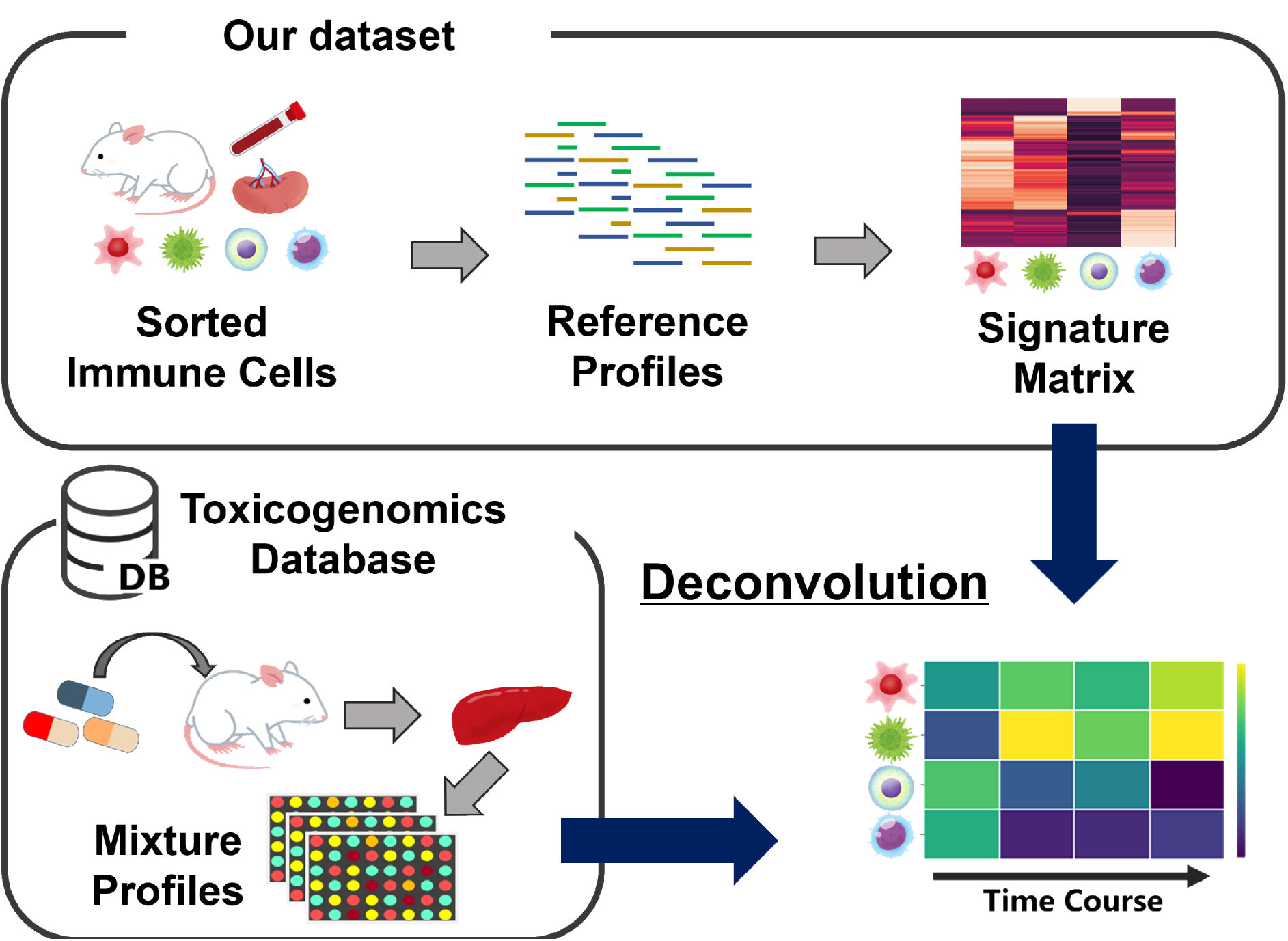
Overview of the present study. The rat-specific ”signature matrix” comprising marker genes that are enriched in each immune cell type was created from our sorted immune cell transcriptome (“reference profiles”). Deconvolution using the prepared signature matrix was applied to the rat liver transcriptome (“mixture profiles”) and estimated immune cell trafficking was subjected to validation and stratification.

## MATERIAL AND METHODS

### Data Preparation

#### Open TG-GATEs dataset

All raw files were downloaded from the home page of Open TG-GATEs (https://toxico.nibiohn.go.jp) (Igarashi *et al*., 2015). Raw signal intensities for each probe set as they are contained in the CEL files, were analyzed using MAS5.0, which is implemented in the software package Bioconductor (http://bioconductor.org) with R (version 3.6.3).

#### Rat RNA sequencing dataset

FASTQ files of the rat RNA sequencing data were prepared according to the RNA isolation and sequencing sections. Quality control of all reads was performed using PINSEQ++ (version 1.2.4) with the indicated parameters (trim_left = 5, trim_tail_right = 5, trim_qual_right = 30, ns_max_n = 20, min_len = 30) (https://peerj.com/preprints/27553/). The expression of transcripts was quantified using Salmon (version 1.6.0) with the indicated parameters (validationMappings, gcBias, seqBias) and a decoy-aware index created using Salmon and Rattus_norvegicus.Rnor_6.0.cdna.all was obtained from EMBL-EBI (https://www.ebi.ac.uk/) (Patro *et al*., 2017). Transcripts per kilobase million data were obtained using tximport, which is implemented in the software package Bioconductor with R (version 4.1.3) from quant.sh files created by Salmon.

### Animals

Male Sprague Dawley rats (6 weeks old) were purchased from CLEA Japan (Tokyo, Japan) for RNA sequencing experiments or Oriental Yeast Co. (Tokyo, Japan) to validate the immune response in the present study. All animals were maintained under standard conditions with a reverse dark-light cycle and were treated humanely. Food and water were available ad libitum. The procedures reported in this article were performed in accordance with the guidelines provided by the Institutional Animal Care Committee (Graduate School of Pharmaceutical Sciences, the University of Tokyo, Tokyo, Japan).

### Rat Liver Injury Model

#### For RNA Sequencing

To induce liver injury, acetaminophen (H0190, Tokyo Chemical Industry Co., Tokyo, Japan) or 4,4′-methylene dianiline (M0220, Tokyo Chemical Industry Co.) was administered at 3 g/kg or 300 mg/kg, respectively by gavage. As a vehicle control group, 0.5 (w/v)% methylcellulose 400 solution (133-17815, Fujifilm Wako Pure Chemical Corporation, Osaka, Japan) was given at 10 µL/g. Only those rats in the experimental groups that were subjected to RNA sequencing had fasted between 21:00 and 9:00, which was the administration time.

#### For Open TG-GATEs Validation Study

To validate the immune responses in the liver, metformin at 1 g/kg by gavage (M2009, Tokyo Chemical Industry Co.), tiopronin at 1 g/kg by gavage (T2614, Tokyo Chemical Industry Co.), galactosamine at 1 g/kg by gavage (G0007, Tokyo Chemical Industry Co.), thioacetamide at 45 mg/kg by gavage (T0187, Tokyo Chemical Industry Co.), colchicine at 15 g/kg by gavage (C0380, Tokyo Chemical Industry Co.), or bortezomib at 0.6 mg/kg by i.v. through a tail vein (B5741, Tokyo Chemical Industry Co.) was administered. As a vehicle control group, 0.5 (w/v)% methylcellulose 400 solution at 10 µL/g by gavage or saline at 2.5 mL/kg by i.v. was given. In this validation experiment, no fasting was performed following the Open TG-GATEs protocol.

### Sample Collection and Preparation

#### Liver

Rats were euthanized and the liver and blood were harvested after compound administration. Briefly, the superior vena cava was clipped using a clamp, and the blood was collected through an inferior vena cava into a 1.5-mL tube containing 1 µL heparin (Yoshindo, Toyama, Japan). The collected blood sample was centrifuged for 15 min at 1,700 g, and the supernatant was subjected to Fuji Drychem NX500sV (Fujifilm Corporation, Tokyo, Japan) to quantify the concentrations of alanine aminotransferase, aspartate aminotransferase, and total bilirubin. The liver perfusion was performed through an inferior vena cava using a plastic cannula-type puncture needle (SR-OT1832C, 18G 32 mm) (Terumo corporation, Tokyo, Japan) and 5 mM HEPES (345-06681, Fujifilm Wako Pure Chemical Corporation) /5 mM EDTA (6381-92-6, Dojindo Laboratories, Kumamoto, Japan) Hanks’ balanced salt solution (17461-05, Nacalai Tesque, Kyoto, Japan). Finally, 1/9 of the liver’s largest lobe or 1/6 of all minced liver was subjected to RNA isolation or flow cytometry, respectively. For flow cytometry, the sample was dissociated using gentleMACS (Miltenyi Biotec, North Rhine-Westphalia, Germany) according to the manufacturer’s instructions. Except where noted, phosphate-buffered saline containing 2% fetal bovine serum was used as “wash buffer” thereafter. After using wash buffer to eliminate hepatocytes, the sample was centrifuged at 50 g at 4 °C for 3 min and the pellets were discarded. To eliminate the hepatic erythrocytes, 5 mL of ACK buffer was added to the sample, and the sample was incubated in a 37 °C water bath for 6 min. ACK buffer was prepared by adding 8,024 mg of NH_4_Cl (A2037, Tokyo Chemical Industry Co.), 10 mg of NHCO_3_ (166-03275, Fujifilm Wako Pure Chemical Corporation), and 3.772 mg of EDTA 2Na2H_2_O (6381-92-6, Dojindo Laboratories) into 1 L of pure water. The sample was washed with the wash buffer three times, and then the sample was subjected to flow cytometry.

#### Blood

An untreated was placed under anesthesia, and blood was collected from the heart by cardiopuncture. The blood was washed once with wash buffer once. After removing the supernatant, 15 mL of ACK buffer was added to the sample and then incubated in a 37 °C water bath for 6 min to eliminate erythrocytes. After incubation, 25 mL of the wash buffer was added to the sample, which was then centrifuged at 400 g at 4 °C for 5 min. The incubation and wash operations were repeated twice. After the last wash, the sample was washed with the wash buffer three times, and then subjected to RNA isolation and flow cytometry or cell sorting.

#### Spleen

An untreated rat was euthanized, and the spleen was collected. Single-cell suspensions from the spleen were obtained by gently forcing the tissues through a 70 µm nylon mesh grid in RPMI-1640 (06261-65, Nacalai Tesque). After washing with the wash buffer, 15 mL of ACK buffer was added into the sample, and then incubated in a 37 °C water bath for 6 min to eliminate the splenic erythrocytes. The sample was washed with the wash buffer three times, and then subjected to RNA isolation and flow cytometry or cell sorting.

### RNA Isolation and RNA Sequencing

Total RNA was prepared using ISOGEN II (311-07361, Nippon Gene Co., Tokyo, Japan) and purified using an RNeasy Plus Mini Kit (74136, Qiagen, Limburg, Netherland) with gDNA elimination using an RNase-Free DNase set (79254, Qiagen) according to the manufacturer’s protocols. RNA was used to prepare RNA-Seq libraries with a NEBNext Ultra ll Directional RNA Library Prep Kit for Illumina (E7760L, New England Biolabs, MA, USA). The libraries were sequenced for pair-end reads using a Novaseq 6000 (Illumina, CA, USA). Sequence reads from each cDNA library were processed as described in the Data Preparation section.

### Flow Cytometry

Flow cytometric analysis was performed with FACSAria III (BD Biosciences, NJ, USA), and data were analyzed with FlowJo software (BD Biosciences). The antibodies used are listed in **Table 2**.

### Cell sorting

Cell sorting was performed with a FACSAria III (BD Biosciences, NJ, USA). CD4 T cells (CD45+/CD11b-/CD3+/CD4+), CD8 T cells (CD45+/CD11b-/CD3+/CD8a+), NK cells (CD45+/CD11b-/CD3-/CD161+), and B cells (CD45+/CD11b-/CD3-/CD45RA+) were obtained from the rat spleen and monocytes (CD45+/CD11b+/CD3-/CD45RA-/CD161-/SSC-low) and neutrophils (CD45+/CD11b+/CD3-/CD45RA-/CD161-/CD43int/SSC-high) were obtained from the rat blood. The antibodies used are listed in **Table 2**. Cells were suspended in wash buffer containing 5% 7-AAD solution (559925, BD Pharmgen) 5 min before analysis to exclude dead cells. The sorted cells were at once pooled in RPMI-1640 containing 10% fetal bovine serum. After centrifugation at 400 g at 4 °C for 5 min, RNA was isolated and sequenced from the cells in the pellets collected.

### Deconvolution

We converted transcript IDs to RGD gene symbols, and median values were selected for duplicate gene names. Only genes in common with the bulk tissue gene expression matrix were retained.

Among the retained genes, up to 200 genes were retained as markers with an absolute fold-change > 1.5 for other cell types. Finally, N genes collected from K cell markers defined above were included in the analysis, and the cell type-specific expression matrix ***X*** ϵ ***R^N×K^*** was defined. Consider a measured bulk gene expression matrix ***Y*** ϵ ***R^N×M^*** for N genes across M samples, each containing K different cell types. The goal of deconvolution is to estimate cell type-specific expression ***X*** ϵ ***R^N×K^*** and cell type-specific proportion ***P*** ϵ ***R^K×M^***, and can be written as:

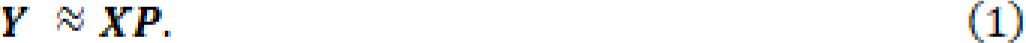

Elastic Net is a regularized regression model with combined L1 and L2 penalties. We can estimate the cell type-specific expression **P̂** via:

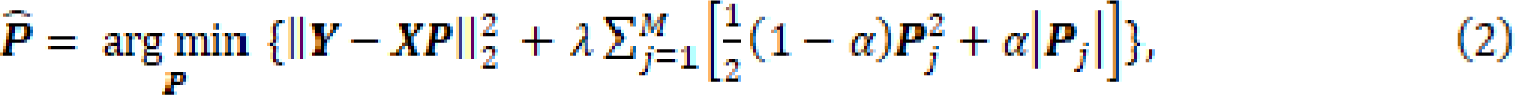

where and are hyperparameters, and we set **λ = 1 and α = 0.05** which are based on published paper using Elastic Net for deconvolution (Altboum *et al*., 2014). The essential codes are available in GitHub (https://github.com/mizuno-group/Deconv). The list of signature genes developed in this study is available in GitHub (RatDeconvolution/result/reference). Analysis was performed using Python (version 3.9.12) with open-source packages (numpy, version 1.22.4; pandas, version 1.1.5; scipy, version 1.8.0; scikit-learn, version 1.0.2; statsmodels, version 0.13.2; umap, 0.5.3)

### Other Species’ Signature Matrices

We used LM6 from the CIBERSORTx home page (https://cibersortx.stanford.edu/) as the human-derived signature matrix (B. Chen *et al*., 2018). For the mouse-derived signature matrix, we collected mouse immune cell transcriptomes from the public database and summarized them to create reference profiles. To convert the gene names, the correspondence tables for rat gene symbols (RGD) to human gene symbols (HGNC) and mouse gene symbols (MGI) were downloaded from the BioMart Ensemble database (http://asia.ensembl.org/index.html). Genes that corresponded to RGD genes that do not exist were omitted from subsequent analysis. The locations and immune cell types of the data are listed in the **Supplementary Data 4**.

### Single Sample Gene Set Enrichment Analysis

The GO annotation file (rgd.gaf) and GO tree structure (go.owl) were downloaded from the GO Consortium website (http://geneontology.org). The ontology at the fifth depth from the top of the tree structure and the corresponding gene set were extracted and used for the following analysis. We subjected RNA sequence data to single-sample gene enrichment analysis (ssGSEA) with following the ssGSEA algorithms (https://github.com/broadinstitute/ssGSEA2.0). Note that the gene expression profiles of the Open-TGGATEs database have approximately three duplicate samples for each administration condition. Figures about the ssGSEA include all those duplicates.

### Clustering Analysis

Dimensionality reduction and clustering were performed using immune cell trafficking or transcriptome of the liver at 3, 6, 9, and 24 h after administration as features. All features were reduced in dimensionality by locally linear embedding, multidimensional scaling, spectral embedding, principal component analysis, t distributed stochastic neighbor embedding, and uniform manifold approximation and projection. The dimensionally reduced features were combined and visualized using a meta-visualization method (Ma *et al*., 2023). The combined meta-distance matrix was subjected to hierarchical clustering and visualized with a heat map. Note that only CD4 T cells, CD8 T cells, monocytes, and neutrophils were used for immune cell trafficking features, and all features were converted to a *z*-score for the corresponding control samples. Additionally, note that transcriptome features were log-transformed and converted to a *z*-score for each compound feature. Adding to immune cell trafficking, some of pickled hematology data (ALT, total bilirubin, platelets, red blood cells, lymphocytes in the blood) were collected from Open TG-GATEs database. All features were converted to a z-score for the corresponding control samples and their time courses were visualized. The detailed code used is found on GitHub (https://github.com/mizuno-group/RatDeconvolution).

Analysis was performed using Python (version 3.9.12) with open-source packages (numpy, version 1.22.4; pandas, version 1.1.5; scipy, version 1.8.0; scikit-learn, version 1.0.2; statsmodels, version 0.13.2; umap, 0.5.3)

## RESULTS

### Extension of Application of Deconvolution to Rats

Deconvolution requires gene expression profiles of the sample to be analyzed (mixture profiles) and gene expression profiles of the target cells with the specific marker genes (signature matrix) created from reference profiles. Given that ***M* ϵ R^n=p^** are the mixture profiles, ***B* ϵ R^k=n^** is the signature matrix, and ***F* ϵ R^n=x^** indicates the scores corresponding to the cell composition, deconvolution calculates F in the following equation: ***M = F = B***, where ***n, p***, and ***k*** are the number of samples, genes, and the estimated cell types.

To extend the application of deconvolution to rats, we obtained gene expression profiles of representative immune cell types for the rat-derived reference profiles. Neutrophils, monocytes, CD4 T cells, CD8 T cells, natural killer cells, and B cells were selected and sorted from the spleen or blood of rats. The immune cell types used in this study are equivalent to LM6, the well-known reference profiles for humans composed of the representative 6 immune cell types prepared by Chen et al. (B. Chen *et al*., 2018). Spleen and blood, whose immune cell composition is known to be different, and the liver derived from rats treated with liver toxicants were obtained and subjected to both transcriptome analysis and flow cytometry simultaneously for validation. As liver toxicants, we selected acetaminophen and methylene dianiline, which are known to induce neutrophil and monocyte migration 24 h after treatment, respectively (Mitchell *et al*., 1973; Bailie *et al*., 1994).

We compared the ratio of immune cells measured by flow cytometry and those estimated by applying deconvolution using the prepared rat-specific reference profiles to the counterpart mixture profiles. Regarding spleen and blood, the results showed that a positive correlation was observed for CD4 T, CD8 T, NK cells, neutrophils, and B cells between the evaluated tissues, whereas the estimate for monocytes varied (**Figure 2A**). The spleen contains resident macrophages with profiles similar those to monocytes (Davies *et al*., 2013; Ingersoll *et al*., 2011), which could interfere with deconvolution and be the cause the variation. We performed the same analysis on liver samples subjected to chemical perturbations and found a high positive correlation, especially for neutrophils and monocytes (**Figure 2B**).

**Figure 2.**
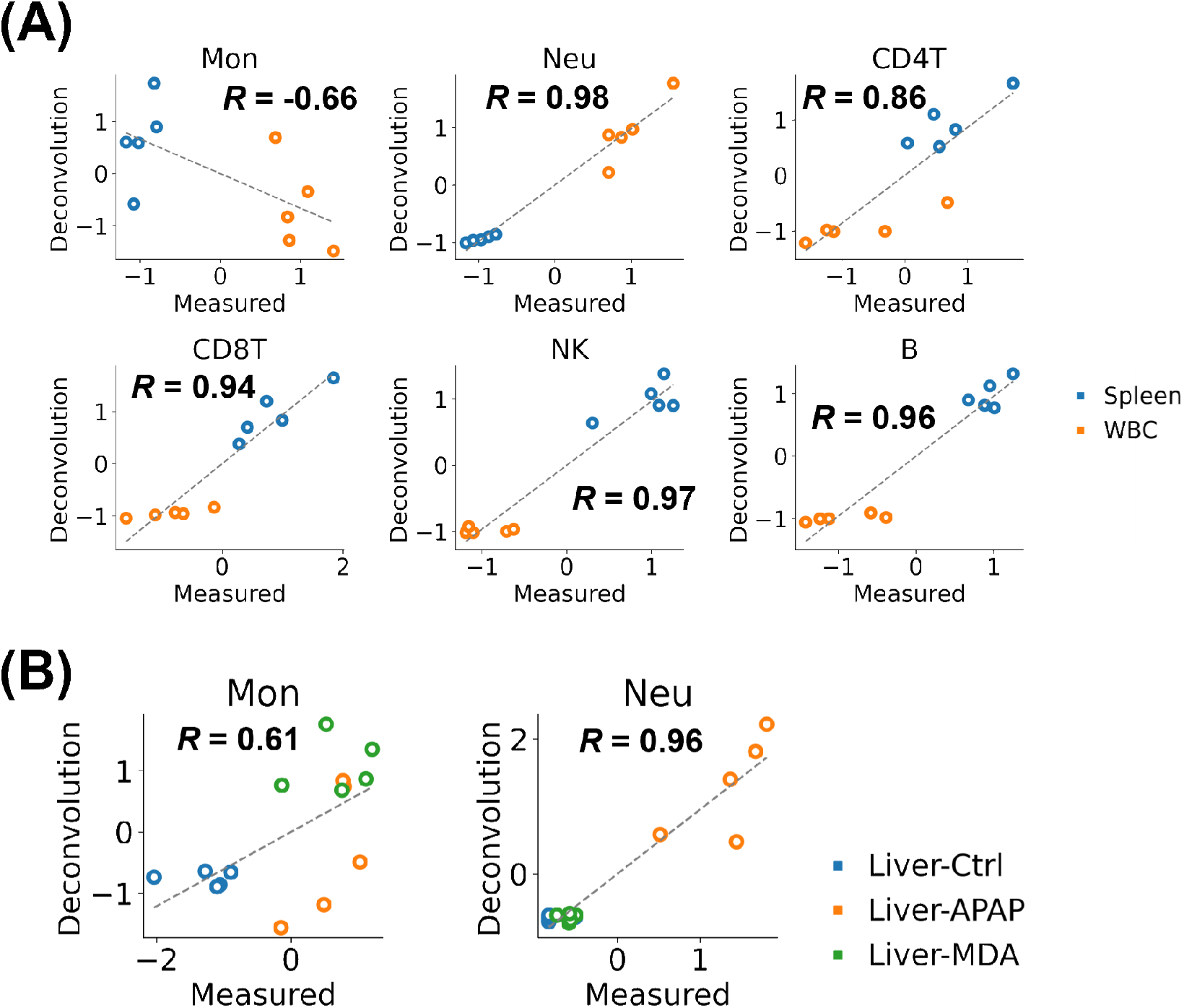
Comparison of estimated and measured immune cell trafficking in rat tissues. (A) Comparison in rat spleen (blue) and whole blood cells (WBC, orange). (B) Comparison using the rat liver after the compounds were administered. Measurement values were obtained with the flow cytometry after cell isolation and red blood cell lysis. Estimated values were calculated by deconvolution using a rat-specific signature matrix. Concordance was measured by Pearson correlation (R) and linear regression (dashed line). Note that these values were converted to z-score between the compared samples. Ctrl, control (blue); APAP, acetaminophen (orange); MDA, methylene dianiline (green).

We also examined the extrapolation using human and mouse signature matrices derived from existing reference profiles. The accuracy was very low especially for CD4 T and CD8 T cells, indicating that there are species differences in the expression of the gene cluster of each reference immune cell and that the rat gene expression profiles are required to create a signature matrix for rat deconvolution (**Supplementary Figures S1 and S2**).

Based on these findings, we concluded that deconvolution in rat gene expression profiles had been established. Notably, in neutrophils and monocytes, the accumulation by perturbation of the compound can be readily estimated with deconvolution. However, care should be taken regarding the limited predictivity and sensitivity of the proposed method. In the comparison analysis of the liver, the estimation accuracy was low for some immune cells including NK cells (**Supplementary Figure S3**). Notably, robust positive correlations were observed across all immune cell types in relation to the spleen. Comparable positive correlations were evident in the blood, with exceptions in the case of monocytes and NK cells (**Supplementary Figure S4**). Conversely, within the liver, we observed low correlations across nearly all immune cell types under a single drug condition. It should be noted that the influence of one outlier sample regarding measurement values obtained by FACS was prominent in the evaluation of neutrophils within the liver control group, as well as NK cells within the liver acetaminophen group, which exhibited low correlation. Combined with the fact that the preparation of liver samples is more complicated than for spleen and blood, it implies that detecting small variances within single condition groups is challenging even when employing experimental methodologies.

Much larger sample size would be necessary for appropriate evaluation of such small variances within a single condition group. In summary, the established deconvolution method exhibits comparable accuracy in estimating immune cell trafficking for inter-condition sample comparisons. However, its efficacy in capturing subtle variances within a single condition group remains uncertain. These findings delineate a limitation of our method and underscore the significance of elucidating the lower threshold of predictable variation for subsequent investigations.

### Validation of Rat-specific Deconvolution to a Public Toxicogenomics Database

Open TG-GATEs is a representative rat toxicogenomics database in which more than 150 compounds were administered to rats at multiple concentrations, and blood biochemical values, liver and kidney gene expression profiles, and liver pathology specimens were obtained at multiple time points (Igarashi *et al*., 2015) . In the present, we used this database for further analysis.

We validated whether the rat-specific deconvolution could be applied to the liver gene expression profiles of Open TG-GATEs, which shows generalized performance of our approach to external and public databases. From the compounds in the database, we targeted compounds causing ALT elevation, which indicates liver toxicity, within 24 h and analyzed these data with the rat-specific deconvolution. Considering that there was no existing information regarding the relationships with the focused immune cells, four compounds were selected as candidates for validation. Colchicine and bortezomib were selected to verify whether the estimated NK and B cell fluctuations could be confirmed. In addition, galactosamine and thioacetamide were selected to compare whether the estimated fluctuations, including those in the early time of 3 h, could be confirmed (**Figure 3A**).

**Figure 3.**
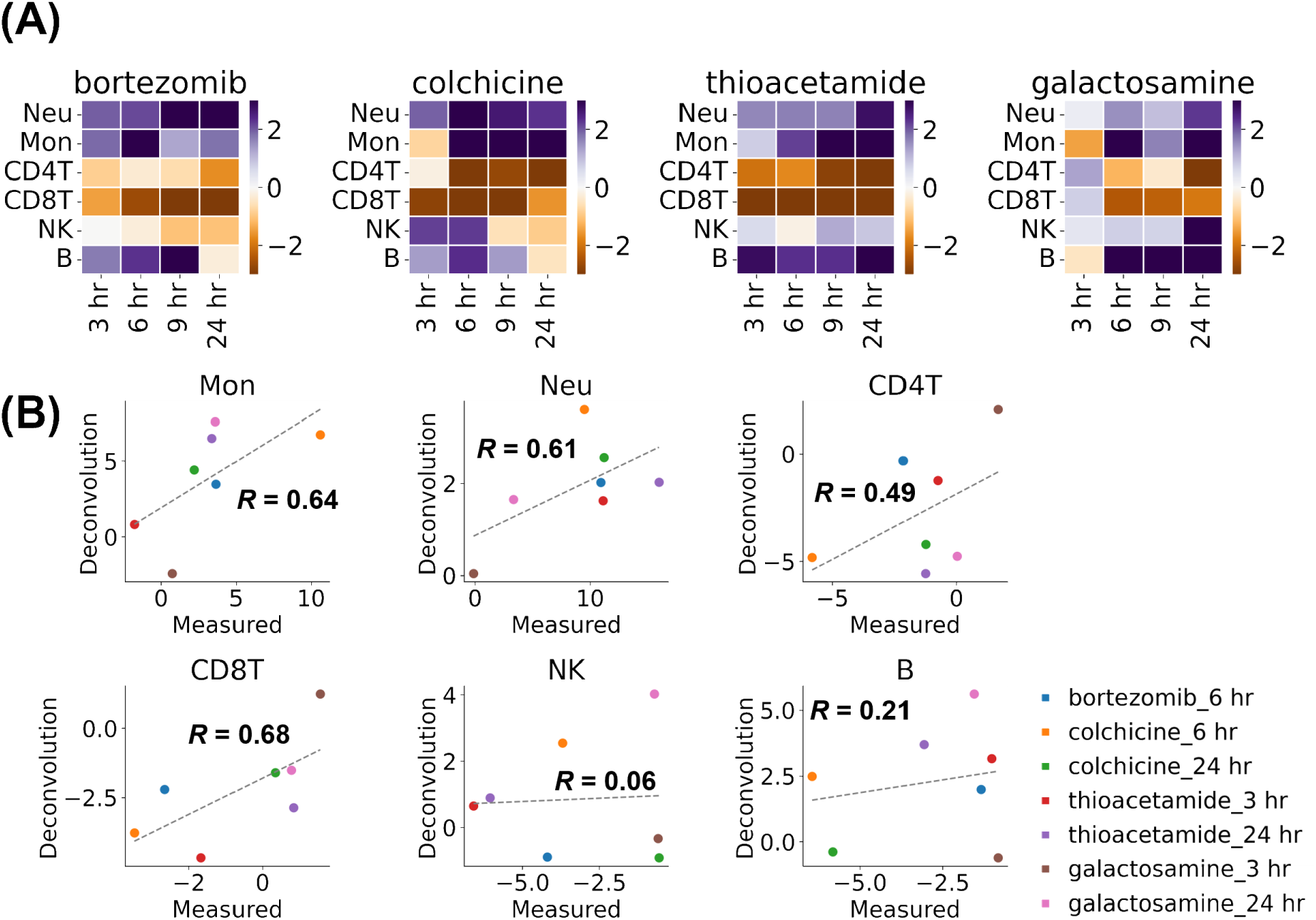
Analysis of Open TG-GATEs database. (A) Heatmaps of estimated immune cell trafficking in the liver. Estimated values at 3, 6, 9, and 24 h after administration of each compound used in the validation are shown. Note that these values are converted to z-score for the corresponding control samples in Open TG-GATEs database or obtained dataset by ourselves. (B) Comparison of estimated and measured immune cell trafficking in rat liver after administering each compound. Concordance was measured by Pearson correlation (R) and linear regression (dashed line). Note that these values are converted to z-scores between the compared samples. Note that each point is the median value in the experiment or in Open-TGGATEs database (n = 3∼4). Mon, monocytes; Neu, neutrophils; CD4T, CD4 T cells; CD8T, CD8 T cells; NK, natural killer cells; B, B cells.

The candidate compounds were administered to rats under the same conditions as those of Open TG-GATEs, and the ratio of immune cells in the liver was measured by flow cytometry. Then, we compared the ratio of immune cells measured by flow cytometry and the estimated values of immune cells by rat-specific deconvolution. Although no clear correlation was observed for NK and B cells, a positive correlation was observed for neutrophils, monocytes, CD4 T cells, and CD8 T cells (**Figure 3B**). It is noteworthy that we confirmed consistency between the observed and estimated results for tiopronin treatment at 3 h: tiopronin induced a weak but significant increase in ALT but no significant change in the neutrophil ratio (**Supplementary Figure S5**). The tiopronin data here indicate that rat-specific deconvolution has the potential to detect these slight differences. One of the explanations for the difficulty in estimating NK and B cells is the possibility of large subpopulations of these cells in the liver. Hepatic NK cells have subpopulations that exhibit different profiles than circulating NK cells in the blood (Mikulak *et al*., 2019). Similarly, B cells have a liver-resident subpopulation that exhibits a specific response during inflammation (Curry *et al*., 2003; Racanelli *et al*., 2001). It is possible that these subsets with close, but different, profiles in the liver, affect the accuracy of deconvolution. Our flow cytometry showed that many NK and B cells were resident in the liver specimens compared with the fewer other immune cells, which may support the impact of liver-resident NK and B cells.

We also examined the extrapolation using human and mouse-derived signature matrices obtained from existing reference profiles. Overall, there were cell types that became less predictable (**Supplementary Figure S5, Supplementary Data 1**). This result supports the previous conclusion and the importance of preparing rat-derived reference profiles for rat deconvolution.

These results suggest that it is possible to estimate and analyze immune cell trafficking except for B and NK cells in the liver from toxicogenomics databases, which are often organized with rats.

### Identification of Clusters of Compounds That Have Different Immune Cell Trafficking

Although there are a few studies on immune cell trafficking as a response to a single compound or a few compounds, there are no reports comparing the responses to a large number of compounds. By applying the established deconvolution to existing large toxicogenomics databases, it is possible to gain aggregated knowledge of the differences in immune cell trafficking as biological responses to variable perturbations. As in the validation study, Open TG-GATEs database was subjected to deconvolution, followed by stratification, and analysis of biological phenomena.

Based on ALT elevation (65), we targeted 16 compounds (**Supplementary Figure S6**, **Table 1**). For the four types of cells (neutrophils, monocytes, CD4 T cells, and CD8 T cells) that had been accurately estimated, as shown in **Figure 3**, the estimated scores of each cell at each time point were used as the features.

**Table 1.** Compound used in stratification analysis. Cluster numbers stratified by analysis are attached for immune-based and transcript-based.

| **compound name** | **immune cluster** | **transcript cluster** |
| --- | --- | --- |
| Tacrine | 4 | 2 |
| Nitrofurazone | 4 | 2 |
| Enalapril | 1 | 2 |
| Metformin | 1 | 4 |
| Gefitinib | 1 | 1 |
| Simvastatin | 1 | 2 |
| Tiopronin | 1 | 2 |
| Naphthyl isothiocyanate | 1 | 1 |
| Bromobenzene | 1 | 3 |
| LPS | 3 | 3 |
| Cycloheximide | 3 | 3 |
| Bortezomib | 2 | 4 |
| Methylene Dianiline | 2 | 4 |
| Galactosamine | 2 | 4 |
| Colchicine | 2 | 3 |
| Thioacetamide | 2 | 4 |

**Table 2.** List of antibodies and staining reagents used in sorting and flow cytometry of immune cells.

| **Marker** | **Detection** | **Cat. No.** | **Conjugate** | **Clone** | **Isotype** |
| --- | --- | --- | --- | --- | --- |
| CD11b | FITC | 561684 | FITC | WT.5 | Mouse IgA, κ |
| CD45 | APC-Cy7 | 561586 | APC-Cy7 | OX-1 | Mouse IgG1, κ |
| CD45RA | BV421 | 740043 | BV421 | OX-33 | Mouse IgG1, κ |
| CD3 | BV605 | 563949 | BV605 | 1F4 | Mouse IgM, κ |
| CD4 | BV786 | 740912 | BV786 | OX-35 | Mouse IgG2a, κ |
| CD8 | BV711 | 740724 | BV711 | OX-8 | Mouse IgG1, κ |
| CD43 | PE | 202812 | PE | W3/13 | Mouse IgG1, κ |
| CD161 | APC | FAB9766R |  |  |  |
| 7AAD | 7-AAD | 559925 |  |  | - |

We used meta-visualization and 6 dimensional reduction methods to visualize and extracted the intrinsic structure of the normalized dataset (**Figure 4A, Supplementary Figure S8**) (Ma *et al*., 2023). Meta-visualization is a spectral method for assessing and combining multiple data visualizations produced by diverse algorithms, generating a consensus visualization that has improved quality over individual visualizations in capturing the underlying structure. This method is theoretically grounded in a general signal-plus-noise model and has robustness against noise and possible adversarial candidate visualizations. The compounds were divided into four clusters based on hierarchical clustering using the combined features by meta-visualization (**Figure 4B**). Note that the result of this clustering is different from the result of clustering of the liver gene expression profiles (**Figures 4C, and 4D, Supplementary Figure S9**). This indicates that the analysis based on immune cell trafficking captures a different perspective from that of gene expression profiles themselves and adds an interpretable new layer.

**Figure 4.**
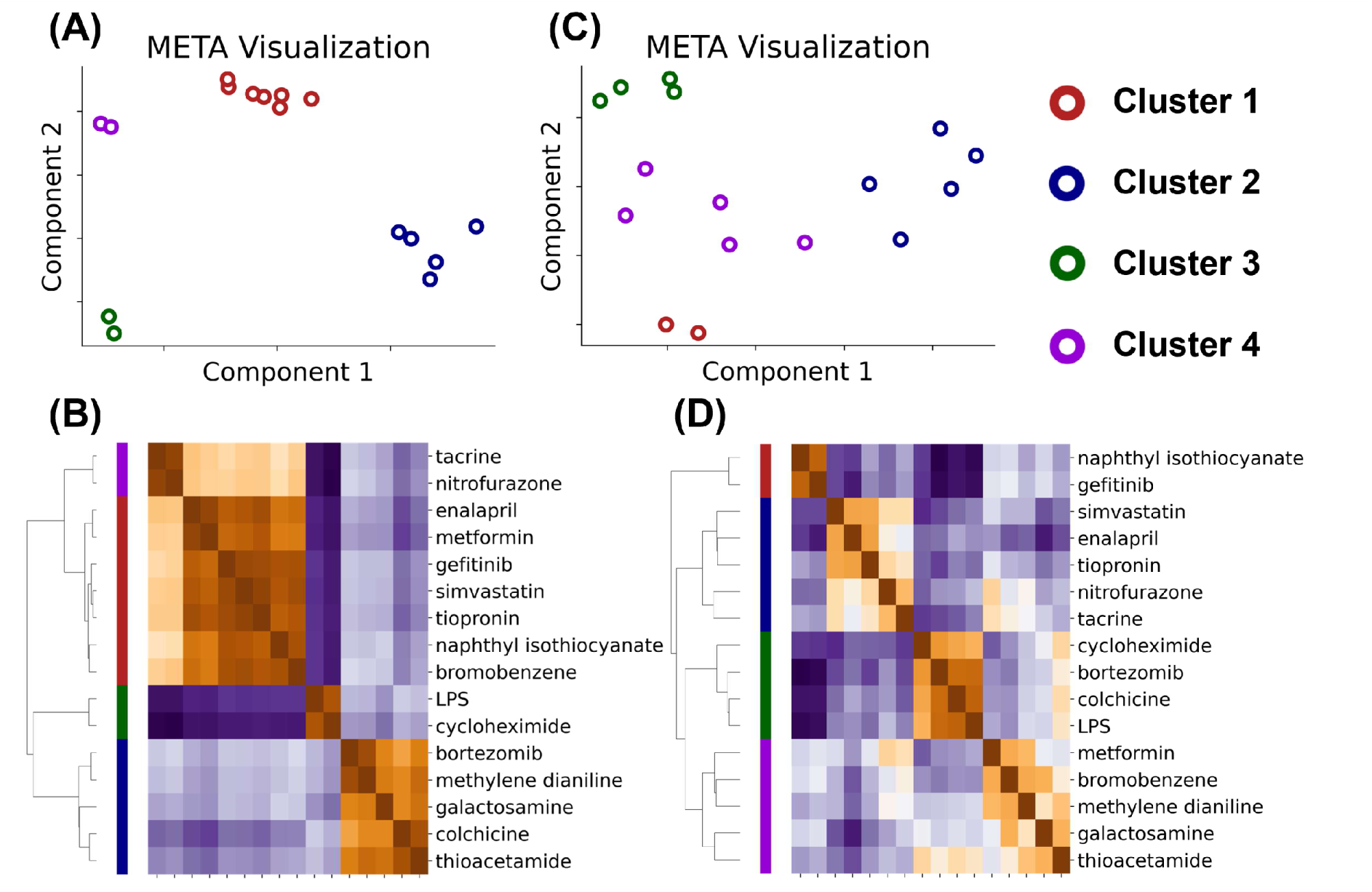
Clustering analysis of 16 compounds that induced significant ALT elevation in the Open TG-GATEs database. Dimensionality reduction and clustering were performed using (A), (B) immune cell trafficking (C), and (D) transcriptome of rat liver at 3, 6, 9, and 24 hours after administration. All features were reduced in dimensionality by locally linear embedding, multidimensional scaling, spectral embedding, principal component analysis, t distributed stochastic neighbor embedding, and uniform manifold approximation and projection. (A), (C) The dimensionally reduced features were combined and visualized by the meta-visualization method. Cluster 1, red; Cluster 2, blue, Cluster 3, green; Cluster 4, violet. (B), (D) The combined meta-distance matrix was subjected to hierarchical clustering and visualized with a heatmap. The color bar closest to the heatmap corresponds to the stratified compound clusters of the heatmap. Note that only CD4 T cells, CD8 T cells, monocytes, and neutrophils were used for immune cell trafficking features. All features were converted to z-score for the corresponding control samples before being subjected to the dimensional reduction methods. Note also that transcriptome features were log-transformed and converted to z-score between each compound feature.

### Characterization of the Compound Clusters

Do the clusters based on immune cell trafficking found in the data-driven analysis in **Figure 4** have any biological meaning? First, to gain insight into the behavior of immune cell trafficking and ALT as the index of liver injury for each cluster, we plotted the time course of their changes (**Figure 5A**). Note that the color filling around the line indicates the 95% confidence interval of all samples belonging to each cluster and depends on the number of compounds. **Figure 5A** shows that compounds belonging to cluster 3 have a strong effect on immune cell trafficking. Cluster 2 is the second strongest cluster on immune cell trafficking, and interestingly, the monocytes at 6 and 9 h after administration show more accumulation than those of cluster 3. Compounds belonging to clusters 1 and 4 have much weaker effects than those of clusters 2 and 3, and cluster 1 is slightly stronger than cluster 4. When these clusters were classified using random forest, ALT did not contribute significantly to the model, suggesting that these clusters do not simply indicate the degree of liver injury (**Supplementary Data 2**). Not surprisingly, because the clusters are different, these behaviors differ from those resulting from clustering based on the transcriptome (**Supplementary Figure S10**).

**Figure 5.**
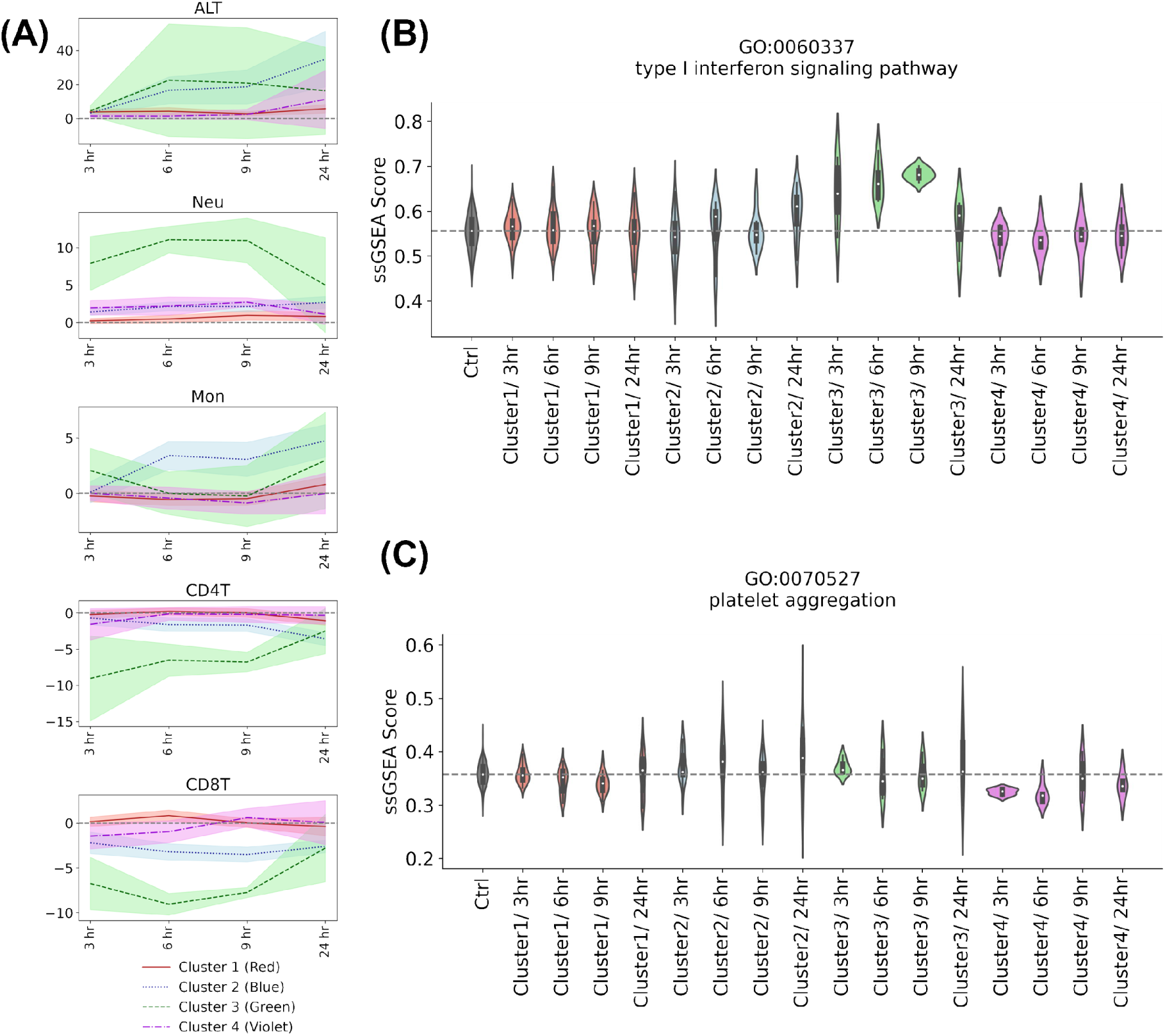
(A) Time course of changes in immune cell trafficking and ALT in stratified compound clusters. The line is the average value for each cluster, and the filled color around it indicates the 95% confidence interval. Each color corresponds to the color of the compound stratified by analysis of immune cell trafficking. Note that the scale of the x-axis does not correspond to the actual time values. (B), (C) Time course of changes in single-sample gene set enrichment analysis (ssGSEA) scores of the liver transcriptome. The ssGSEA scores of gene ontology terms GO: 0060337 type I interferon signaling pathway and GO: 0070527 platelet aggregation were visualized as violin plots. Note that other terms were visualized in Supplementary Figure 12.

Cluster 3, the strongest cluster on immune cell trafficking, contains two compounds, LPS, which is known as a strong inflammation inducer, and cycloheximide, a protein synthesis inhibitor targeting the translational process (Ferluga and Allison, 1978; Ledda-Columbano *et al*., 1992). Because the number of compounds in this cluster is small and easy to compare with existing knowledge, we focused on this cluster and evaluated its consistency with the existing biological knowledge. We calculated the gene ontology enrichment score back to the gene expression data of each compound to investigate the commonalities among the effects of the compounds. We searched for gene ontology terms that discriminate these compounds from those belonging to other clusters with statistical tests. As a result, the type I interferon signaling pathway was detected as a characteristic gene ontology term (**Figure 5B**). Although existing knowledge of the inflammatory response induced by cycloheximide is limited compared to that of LPS, a representative inflammation inducer, in a literature survey, cycloheximide was reported to stimulate the production of type I interferon RNA (Ringold *et al*., 1984).

Similarly, in cluster 4, to which tacrine and nitrofurazone belong, platelet aggregation was detected as a characteristic gene ontology term, particularly in the early time points (**Figure 5C**). Platelets are known to release signaling factors related to liver injury and regeneration, and their association with acute liver failure and liver disease has been reported (Morris and Chauhan, 2022). Their importance has also been suggested in mouse models of drug-induced liver injury (Chauhan *et al*., 2020; Miyakawa *et al*., 2015). A literature survey indicated that the two compounds inhibit platelet aggregation (Rossi and Levin, 1973; Slevin *et al*., 2018). In addition, the biochemical values of blood in the database showed a slight trend toward an increase in platelet concentration at 9 h compared with the other clusters (**Supplementary Figure S11**). The two compounds can influence liver injury by affecting pathways and mechanisms related to platelets.

Other characteristic other gene ontology terms for each cluster were found by statistical tests with conservative correction for multiplicity (**Supplementary Figure S12**, **Supplementary Data 3**).

Biological knowledge, extracted by a stratification analysis using estimated immune cell trafficking, could not be extracted by using the liver transcriptome alone (**Supplementary Figures S13 and S14**). In summary, the clusters obtained by immune cell trafficking are expected to have biological meanings that are consistent with existing biological knowledge, rather than overfitting the data.

## DISCUSSION

### Importance of establishing deconvolution method in rats

Although there have been studies that applied deconvolution to rat tissues, to our knowledge, there are no rat immune cell transcriptome datasets appropriate for reference profiles of deconvolution in public databases, and therefore, LM22, a human dataset, is often employed (Wang *et al*., 2021; Gil Del Alcazar *et al*., 2022). Here, we have confirmed that the use of mouse or human-derived reference profiles for deconvolution of rat specimens resulted in a decrease in accuracy for several immune cell types compared with rat-derived reference profiles, and that species differences did exist. This indicates that the rat-derived reference profiles obtained in this research are necessary for accurate deconvolution of rat specimens. Note that single-cell RNA-seq data of rat specimens, including rat liver, are currently available (Qaisar *et al*., 2021) . However, the characteristics of the data, which would affect the accuracy of the deconvolution, are different from the majority of accumulated legacy transcriptome datasets in public databases.

Deconvolution can now be applied to rats, for which large-scale toxicogenomics data are available in public databases, and work as a knowledge miner for immune cell trafficking in response to various perturbations achieved by compounds. In the present study, we tested data obtained from Open TG-GATEs because of its rich time-series data, while analysis of data from DrugMatrix, another large toxicogenomics database containing a larger number of compounds than Open TG-GATEs, is also an interesting target. Other individual data in the toxicological database, not used in this study, would be interesting as well. Pathological information is one of the most interesting targets for further analysis. Linking immune cell trafficking or compounds cluster to pathology holds the potential to promote our understanding of toxicological mechanisms and prognosis. By aggregating extracted knowledge, we could deepen our understanding of immune cell trafficking. Notably, rats are often used in the field of toxicology such as safety assessment of drugs and chemicals. Thus, the combination of rat-specific deconvolution and toxicity databases would contribute not only to understanding the mechanisms of immune cell trafficking in liver injury but also to toxicology such as stratification of the types of toxicity based on this new layer. Moreover, such efforts will lead to the expansion of organ responses in Adverse Outcome Pathway (Ankley *et al*., 2010).

### Types of immune cells that can be estimated in rat deconvolution

One of the limitations of this study is the restricted number of immune cell types that can be estimated for the experimental and analytical reasons. First, knowledge of rat immunology is much lower than that of mice. The established isolation protocols and the publications for rat immune-relative cells are limited based on our preliminary literature survey. This is an interesting contrast to the field of toxicology, where rats are more often employed than mice. Second, as shown in **Figure 3**, it is difficult to estimate NK and B cells when rat-specific deconvolution is applied to the external data. In reference-based deconvolution, the signature matrix is created so that the co-correlation among immune cells to be analyzed is low. Because the main algorithm of deconvolution in this study is linear regression, model instability is inevitable in the presence of multicollinearity (Dormann *et al*., 2013). A possible solution to this problem is to incorporate the other cells belonging to the tissue of interest in the model. Notably, the importance of modelling considering tissue specificity in the deconvolution method has been discussed, although it has not been rigorously demonstrated. In this regard, the dataset we provide through this study could act as a dataset to evaluate deconvolution on liver tissue.

To increase the number of immune cell types for rat deconvolution, the utilization of scRNA-seq data is one idea (Schelker *et al*., 2017). However, care should be taken in using of scRNA-seq data for reference-based deconvolution because the shape of the scRNA-seq data differs from that of bulk transcriptome data accumulated in public databases. In this regard, unsupervised and semi-supervised deconvolution methods, which are independent or less dependent on the reference profile, respectively, and have been developed in recent years, are good options (Gaujoux and Seoighe, 2012; Dimitrakopoulou *et al*., 2018; Li and Wu, 2019; Tang *et al*., 2020). Comparison of deconvolution method paradigms is a future work for utilizing large toxicogenomics databases as data sources for immune cell trafficking (Avila Cobos *et al*., 2020).

### Contamination of blood into the liver transcriptome

Blood contamination is one of the important discussion topics in the utilization of deconvolution for knowledge miner for immune cell trafficking from public databases. We measured the immune responses and transcriptomes using samples collected after liver perfusion. By contrast, the transcriptomes in the database were only collected after blood was released from the inferior vena cava. This may have resulted in contamination of the samples with blood-derived immune cells, and these differences may have affected the accuracy comparisons. In fact, in some samples, the number of neutrophils in the blood in the database is proportional to the score estimated by deconvolution (**Supplementary Figure S15**). However, it is difficult to determine whether this is the actual behavior of immune cells in the liver or a significant blood influence.

How can immune cells infiltrating tissues and blood-derived immune cells be distinguished and evaluated? Intravascular staining method is a possible experimental solution to this problem (Anderson *et al*., 2014). In this method, staining discriminates between tissue-localized cells and blood-born cells by intravenous injection of staining antibodies. However, the liver differs from organs such as the lungs, where the intravascular staining method is generally applied, in that it has fenestrae, and care should be taken when applying intravascular staining method (Gracia-Sancho *et al*., 2021). Anderson et al referred to the specific character of CD69+ cells regarding the relationship between the liver and blood. In addition, there are few reports about applying intravascular staining method in studies of tissue injury (Anderson *et al*., 2014). The vascular wall discriminating hepatocytes and blood flow is disrupted in liver injury, and whether intravascular staining method can be applied to investigate liver injury remains to be determined (DeLeve *et al*., 1999). It is important to grasp the biological background of the source specimens when using legacy data because it directly affects the interpretation of the data analysis outcomes. Thus, it is essential to understand the relationship between the immune cells infiltrating in the liver and those from blood with a liver injury background.

## DATA AVAILABILITY

RNA-sequencing data, meta information and raw FASTQ files are shown in GEO datasets under accession number GSE239996.

## ACKNOWLEDGEMENTS

We appreciate Dr. Hiroyuki Kusuhara and Dr. Hisamitsu Hayashi for helpful discussion about this study. We thank all those who contributed to the construction of the datasets employed in the present study such as Open TG-GATEs.

## AUTHOR CONTRIBUTIONS

Katsuhisa Morita: Data curation, Formal analysis, Methodology, Software, Investigation, Writing – Original Draft, Visualization.

Tadahaya Mizuno: Conceptualization, Data curation, Investigation, Resources, Supervision, Project administration, Writing – Original Draft, Writing – Review & Editing, Funding acquisition.

Iori Azuma: Data curation, Investigation.

Hiroyuki Kusuhara: Writing – Review

---

**Supplementary Figure S1.**
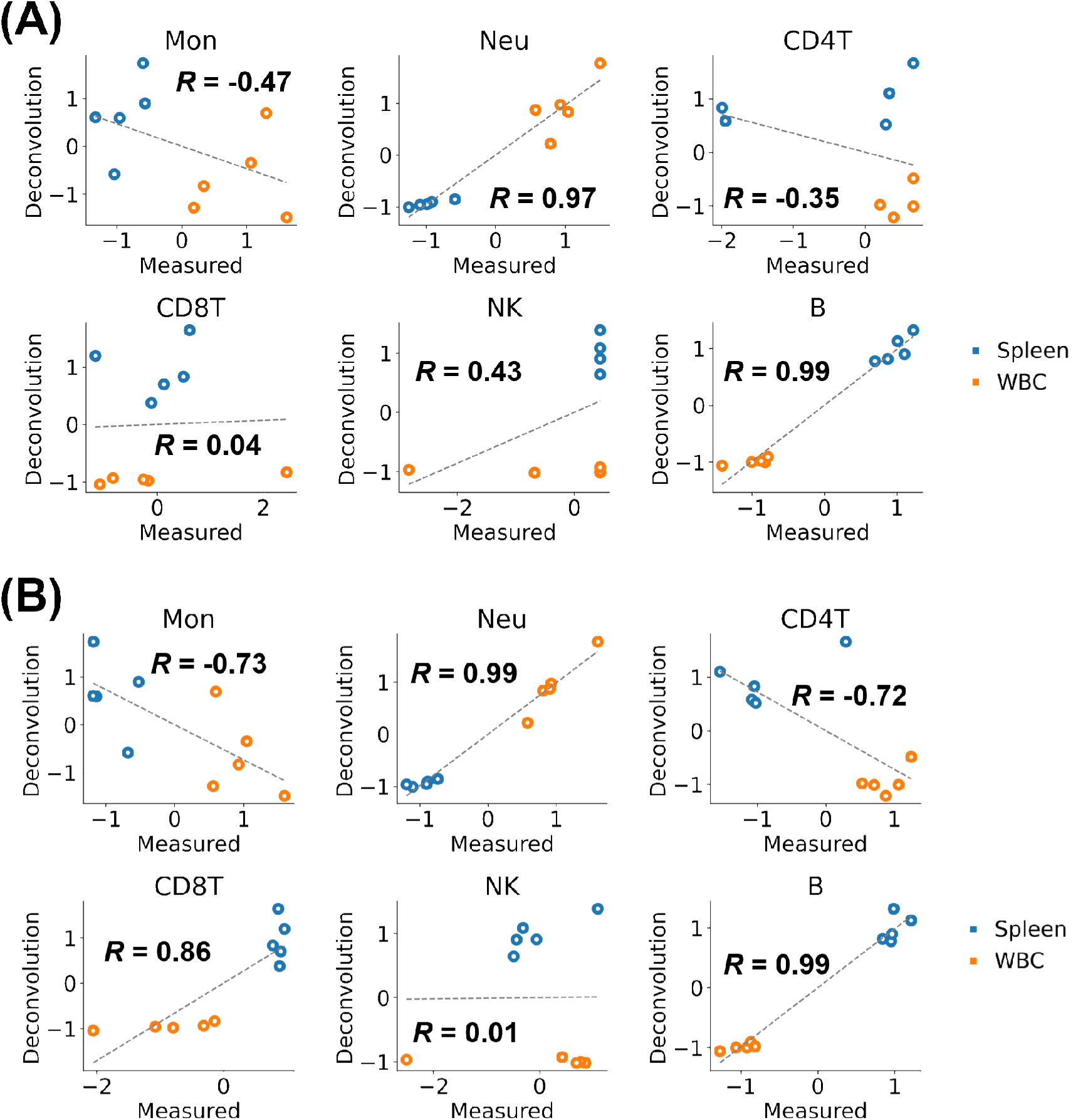
Comparison of estimated and measured immune cell trafficking in the rat spleen and whole blood cells (WBCs). Values were measured by flow cytometry after cell isolation and red blood cell lysis. Estimated values were calculated by deconvolution using (A) a human-derived signature or (B) mouse-derived signature matrices. Note that these values were converted to *z*-score between the compared samples.

**Supplementary Figure S2.**
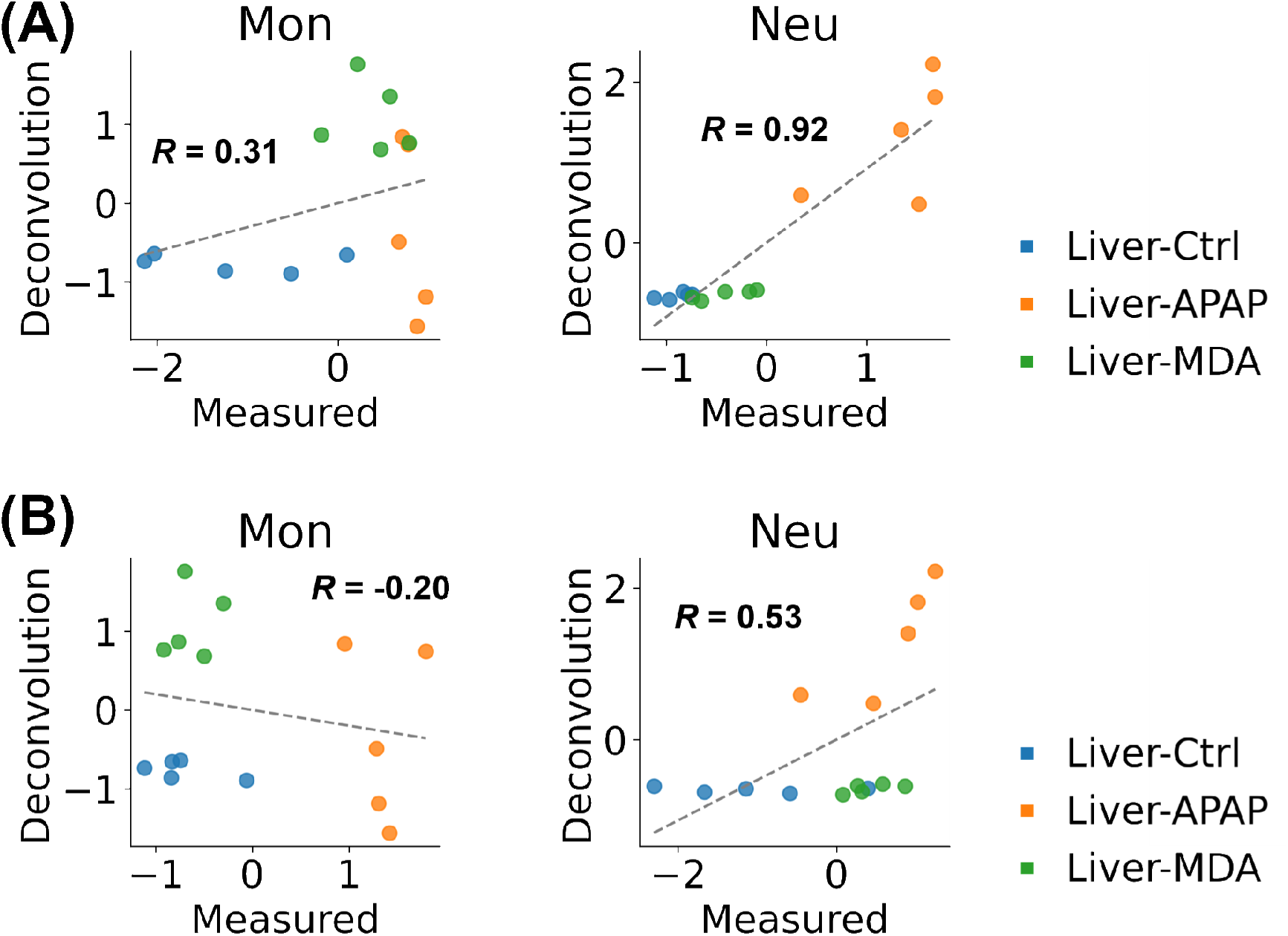
Comparison of estimated and measured immune cell trafficking in rat liver after compounds were administered. values were measured by FACS after cell isolation and red blood cell lysis. Estimated values were calculated by deconvolution using (A) a human-derived signature matrix or (B) mouse-derived signature matrices. Note that these values were converted to *z*-score between the compared samples. Ctrl, control; APAP, acetaminophen; MDA, methylene dianiline.

**Supplementary Figure S3.**
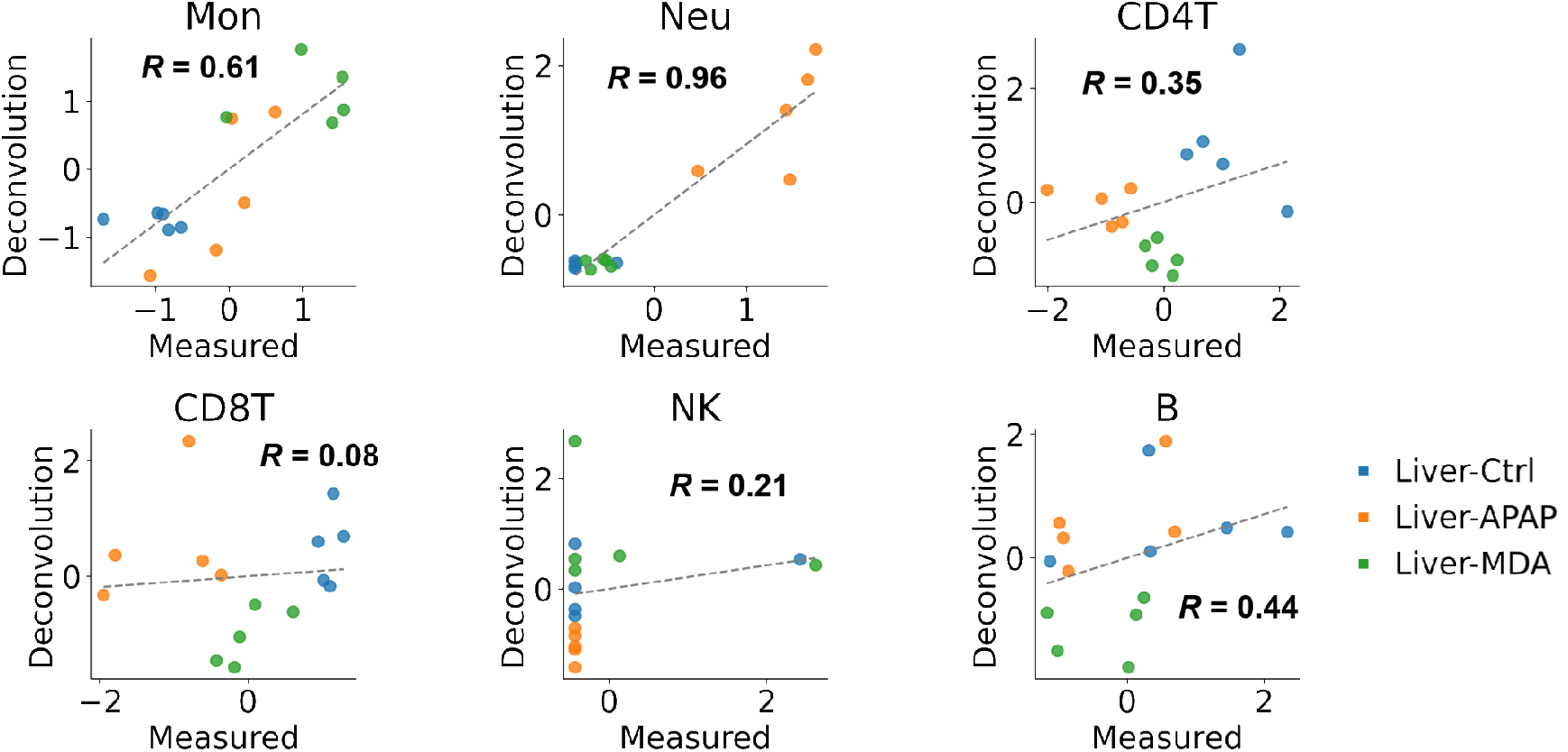
Comparison of estimated and measured immune cell trafficking in rat liver after compounds were administered. Values were measured by FACS after cell isolation and red blood cell lysis. Estimated values were calculated by deconvolution using a rat-specific signature matrix. Note that these values were converted to *z*-score between the compared samples. Ctrl, control; APAP, acetaminophen; MDA, methylene dianiline.

**Supplementary Figure S4.**
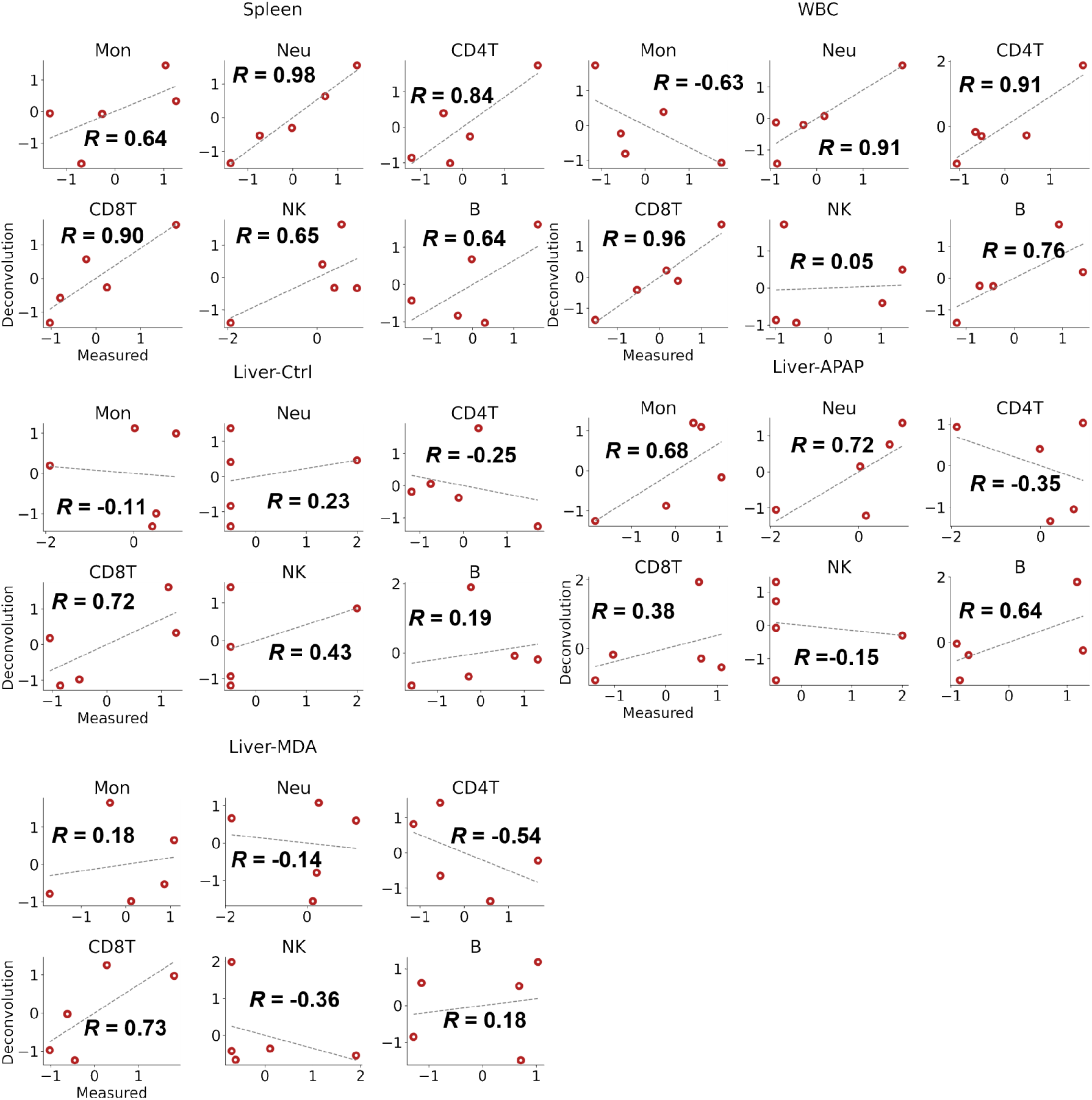
Comparison of estimated and measured immune cell trafficking in the spleen, the blood or the liver after each compound were administered. Measurement values were measured by the flow cytometry after cell isolation and red blood cell lysis. Estimated values were calculated by deconvolution using a rat-specific signature matrix. Concordance was measured by Pearson correlation (R) and linear regression (dashed line). Note that these values were converted to z-score between the compared samples. Ctrl, control; APAP, acetaminophen; MDA, methylene dianiline.

**Supplementary Figure S5.**
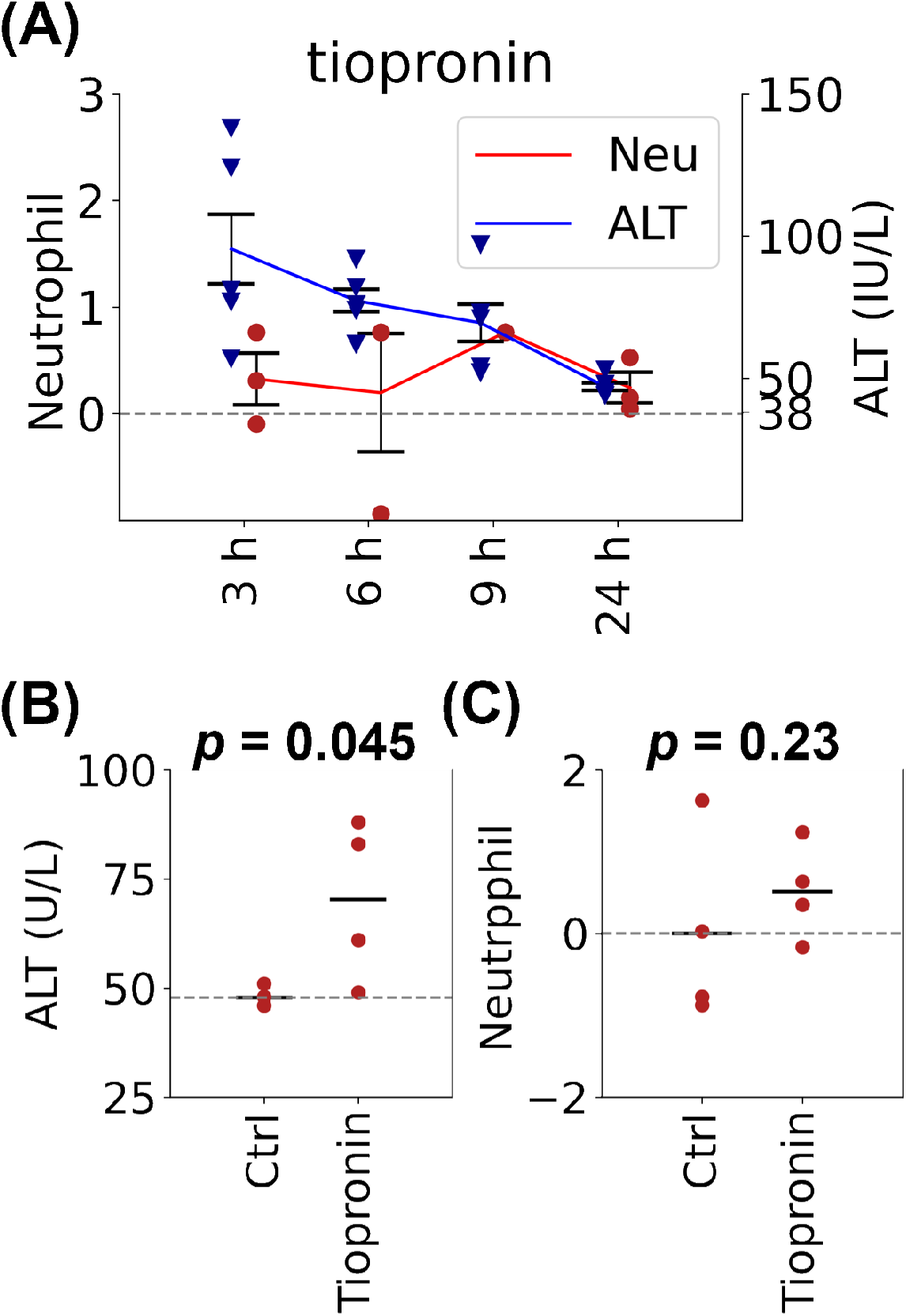
(A) Time course of changes in neutrophils estimated in the liver and ALT measured in the blood of rat administered tiopronin as determined from the Open TG-GATEs database. Value of (B) ALT in blood and (C) neutrophils in the liver at 3 h after tiopronin administration. Note that neutrophils values are converted to *z*-score for the corresponding control samples. The tests of significance were conducted using an unpaired sample greater-sided Welch *t* test.

**Supplementary Figure S6.**
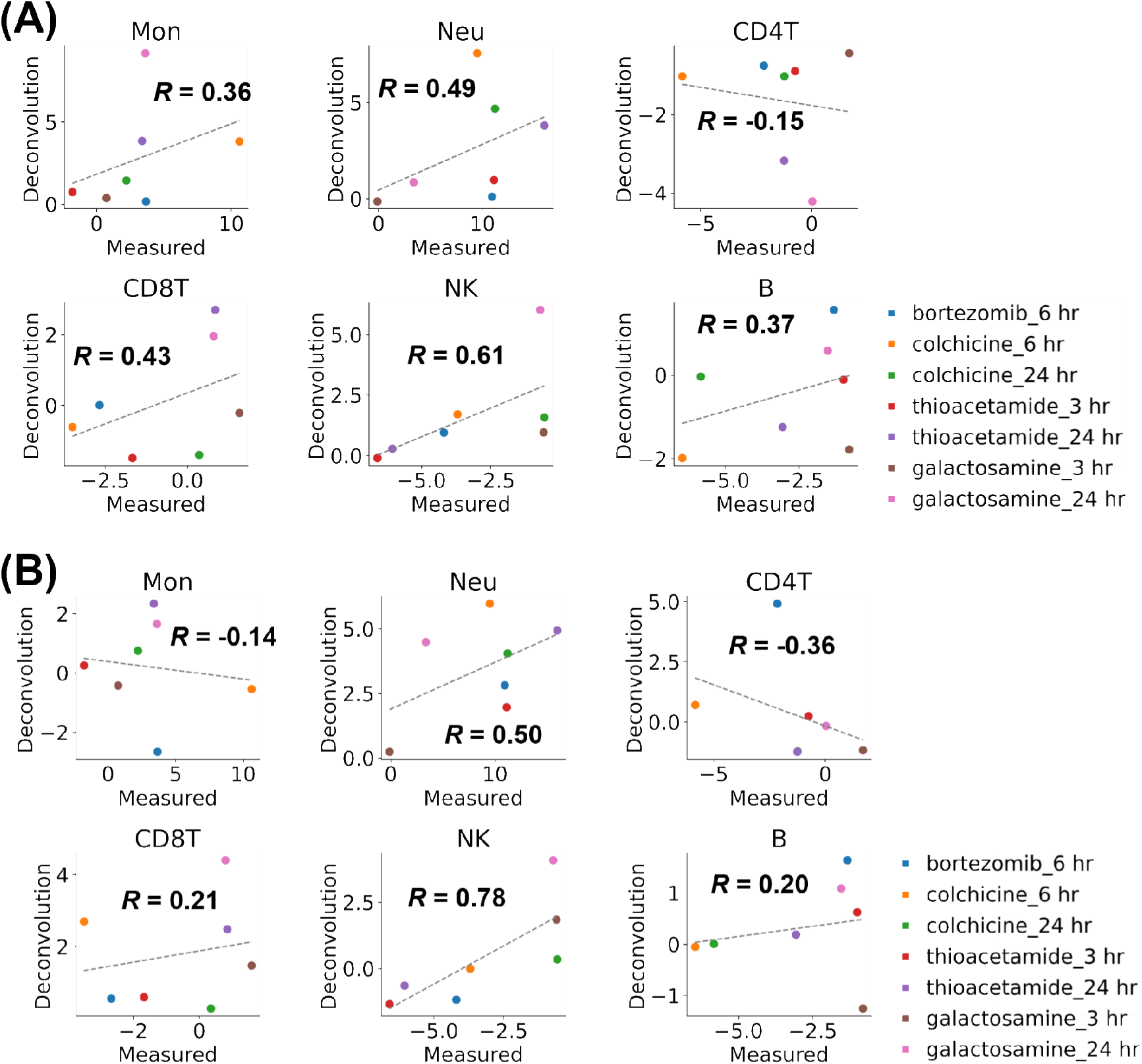
Comparison of estimated and measured immune cell trafficking in the rat liver after administering each compound. Values were calculated by deconvolution using (A) human-derived signature matrix or (B) mouse-derived signature matrices. Note that these values were converted to *z*-score between the compared samples. Note that each point is the median value in the experiment or in the Open-TGGATEs database (n = 3∼4).

**Supplementary Figure S7.**
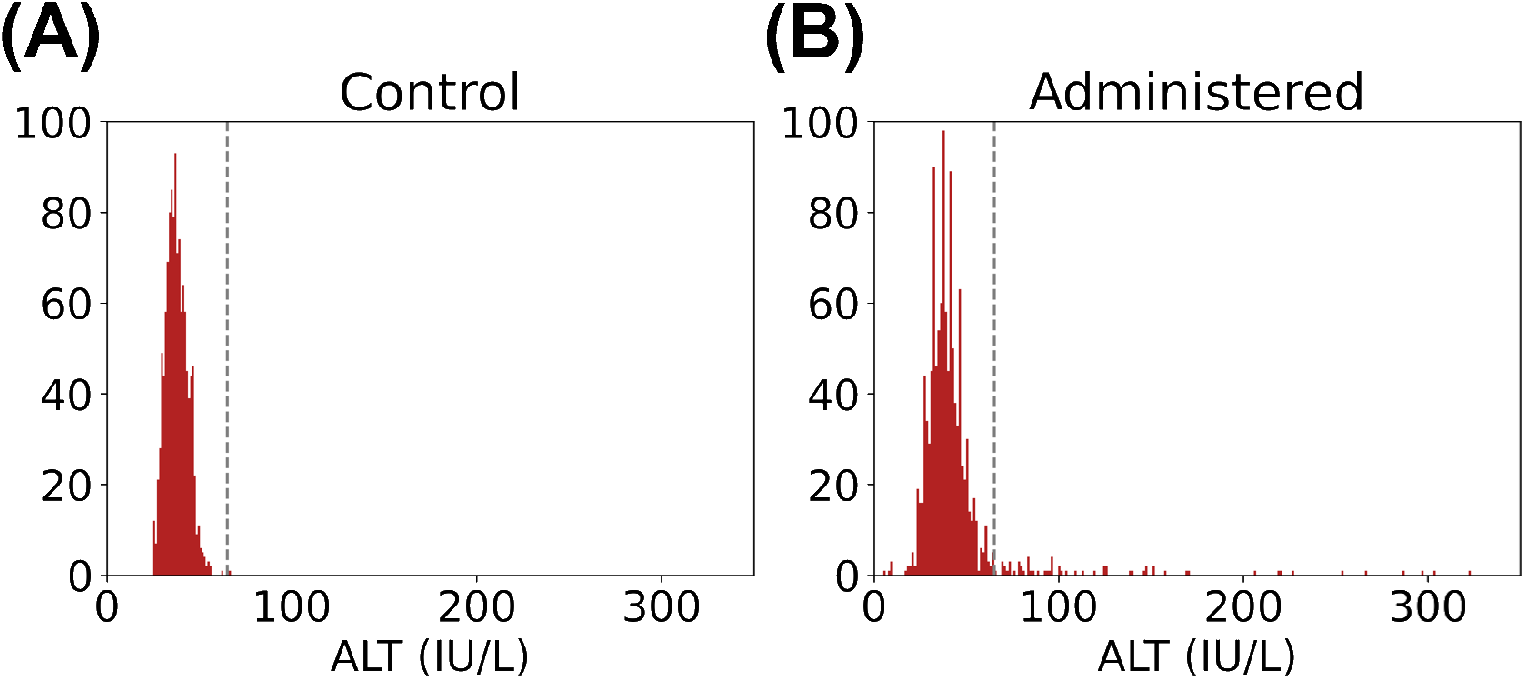
ALT distribution after (A) administration of control solvent or (B) of compound from the Open TG-GATEs database. The vertical dotted line is the threshold for targeting the compounds used for stratification.

**Supplementary Figure S8.**
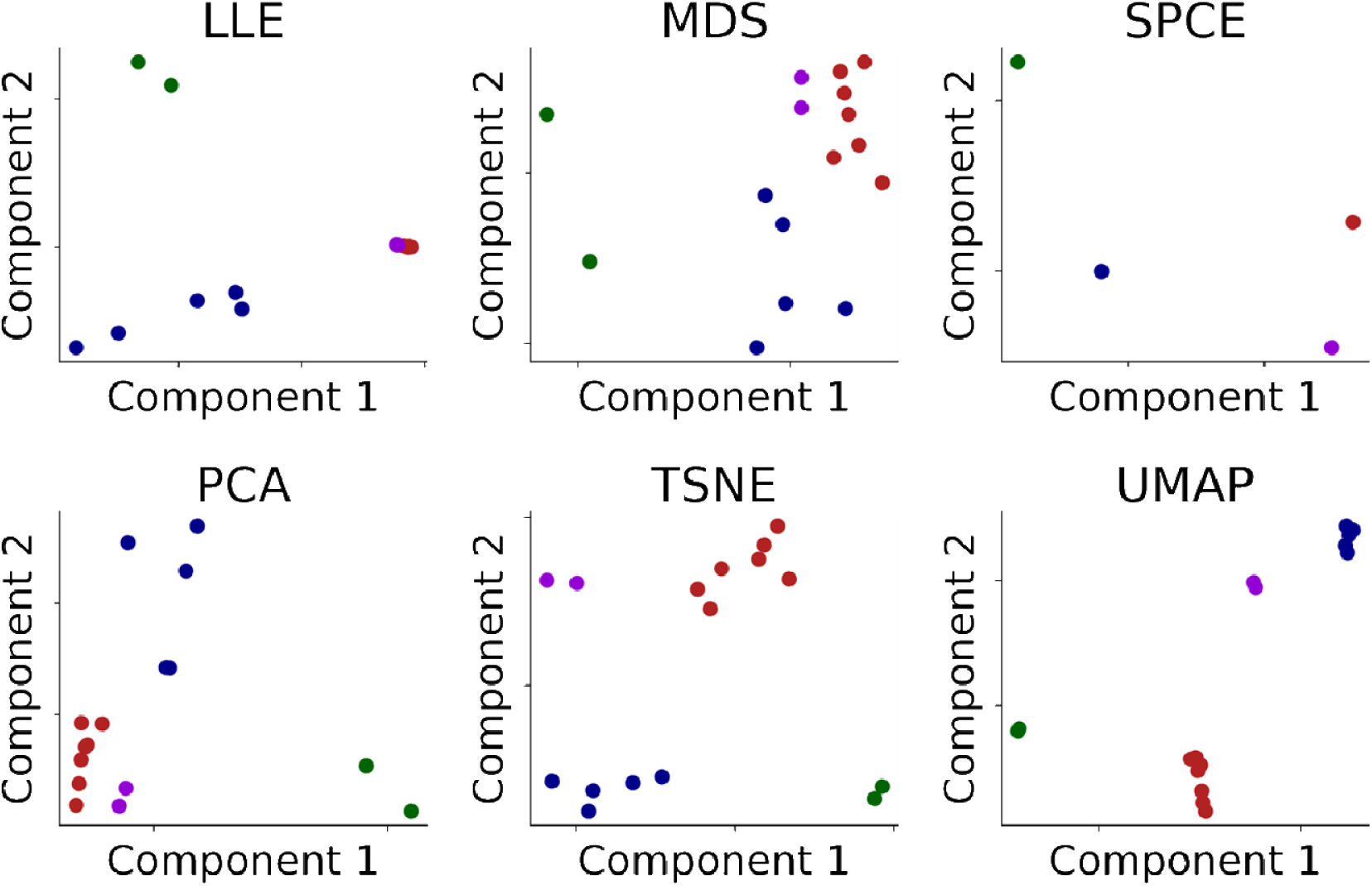
Visualization of immune cell trafficking features of Open TG-GATEs compounds. All features were dimensionally reduced by each method and visualized. These reduced features were subjected to meta-visualization for Figure 4A. Each color corresponds to the color of the stratified compound. LLE, locally linear embedding; MDS, multidimensional scaling; SPCE, spectral embedding; PCA, principal component analysis; TSNE, *t* distributed stochastic neighbor embedding; UMAP, uniform manifold approximation and projection.

**Supplementary Figure S9.**
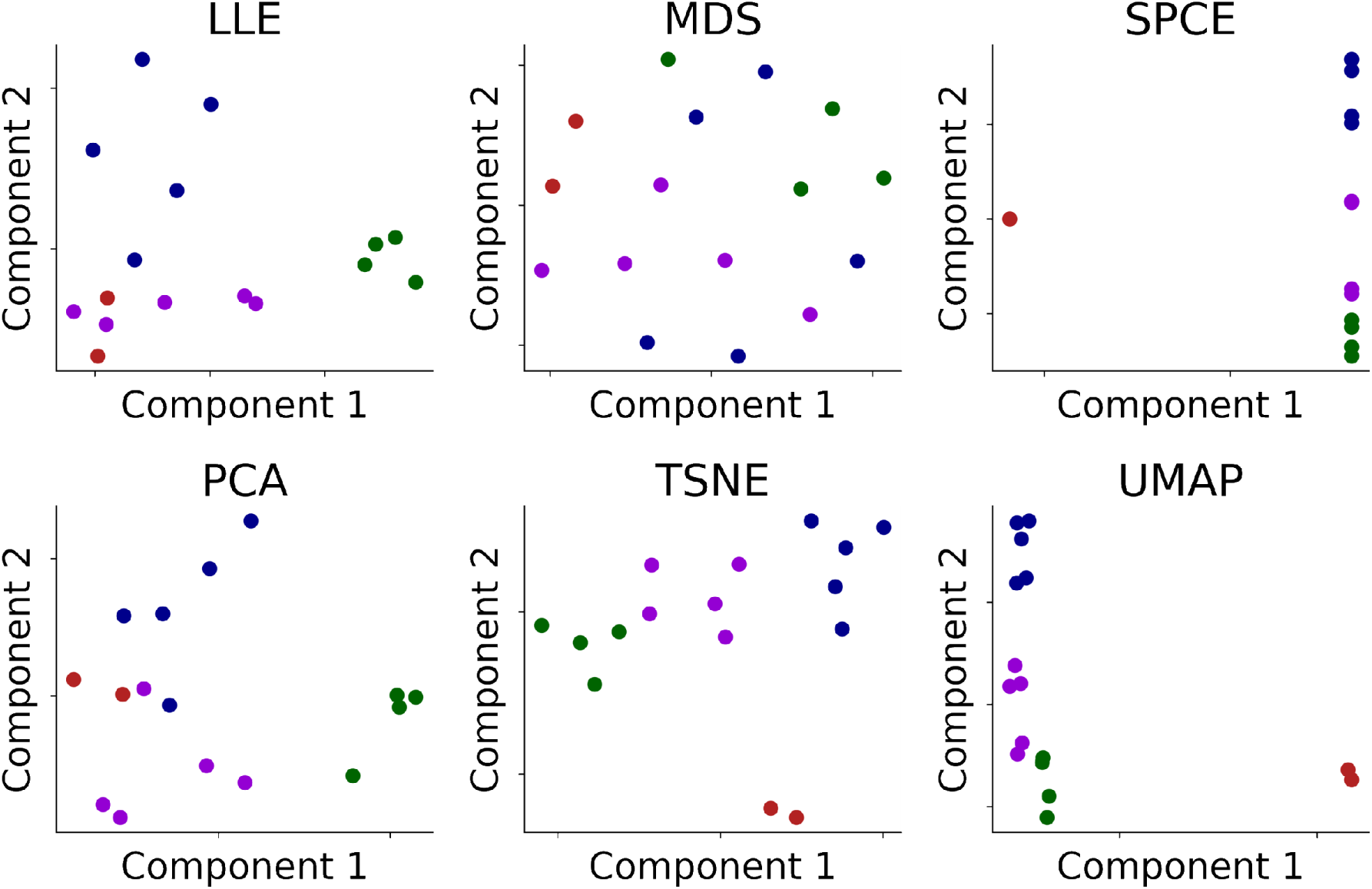
Visualization of transcriptome features of Open TG-GATEs compounds. All features were dimensionally reduced by each method and visualized. These reduced features were subjected to meta-visualization for Figure 4C. Each color corresponds to the color of the compounds stratified by transcriptome analysis. LLE, locally linear embedding; MDS, multidimensional scaling; SPCE, spectral embedding; PCA, principal component analysis; TSNE, *t* distributed stochastic neighbor embedding; UMAP, uniform manifold approximation and projection.

**Supplementary Figure S10.**
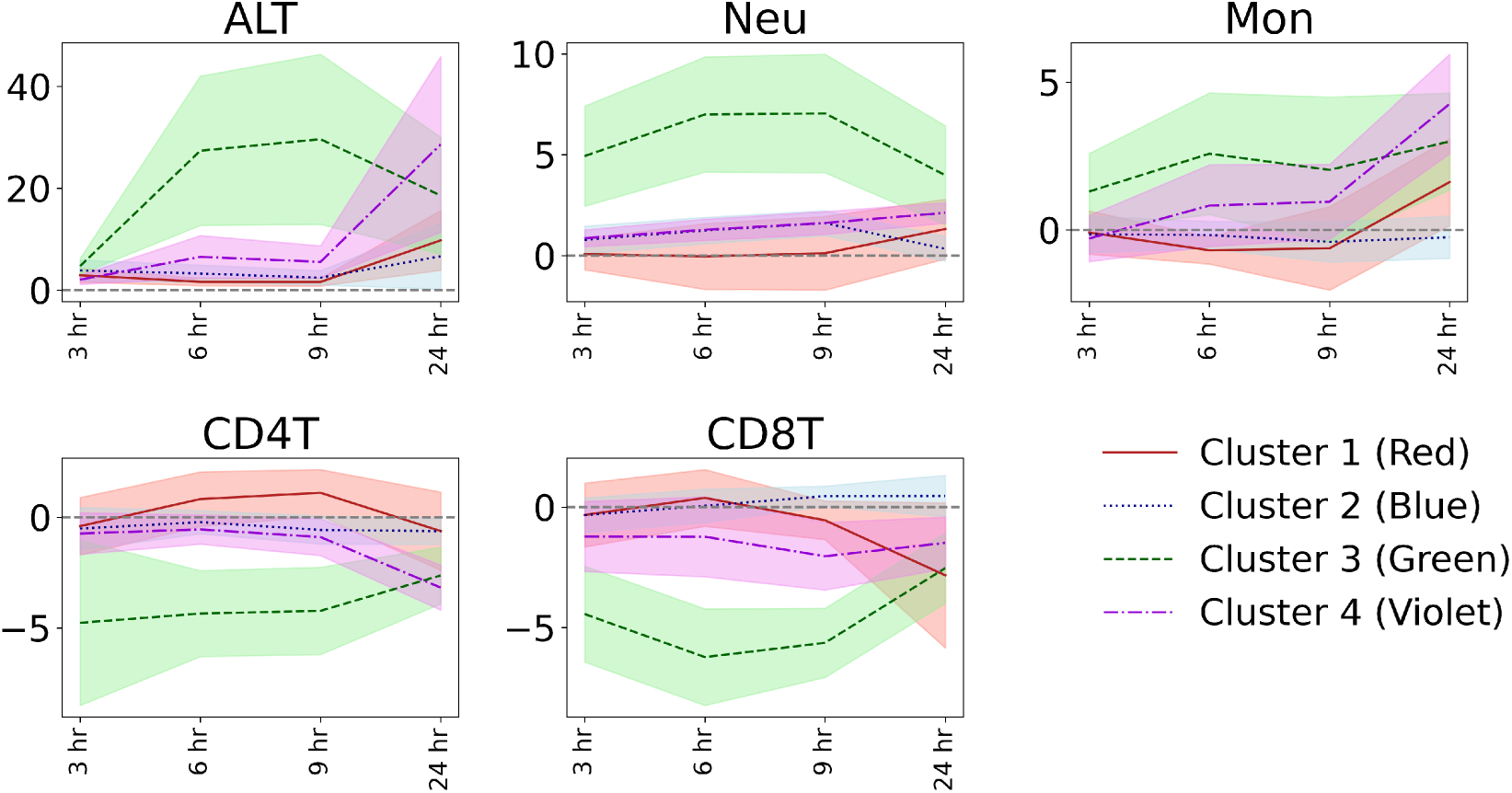
Time course of changes in immune cell trafficking and ALT in stratified compound clusters. The line is the average value for each cluster, and the filled color around it indicates the 95% confidence interval. Each color corresponds to the color of the compound stratified by analysis of the transcriptome. Note that the scale of the x-axis does not correspond to the actual times.

**Supplementary Figure S11.**
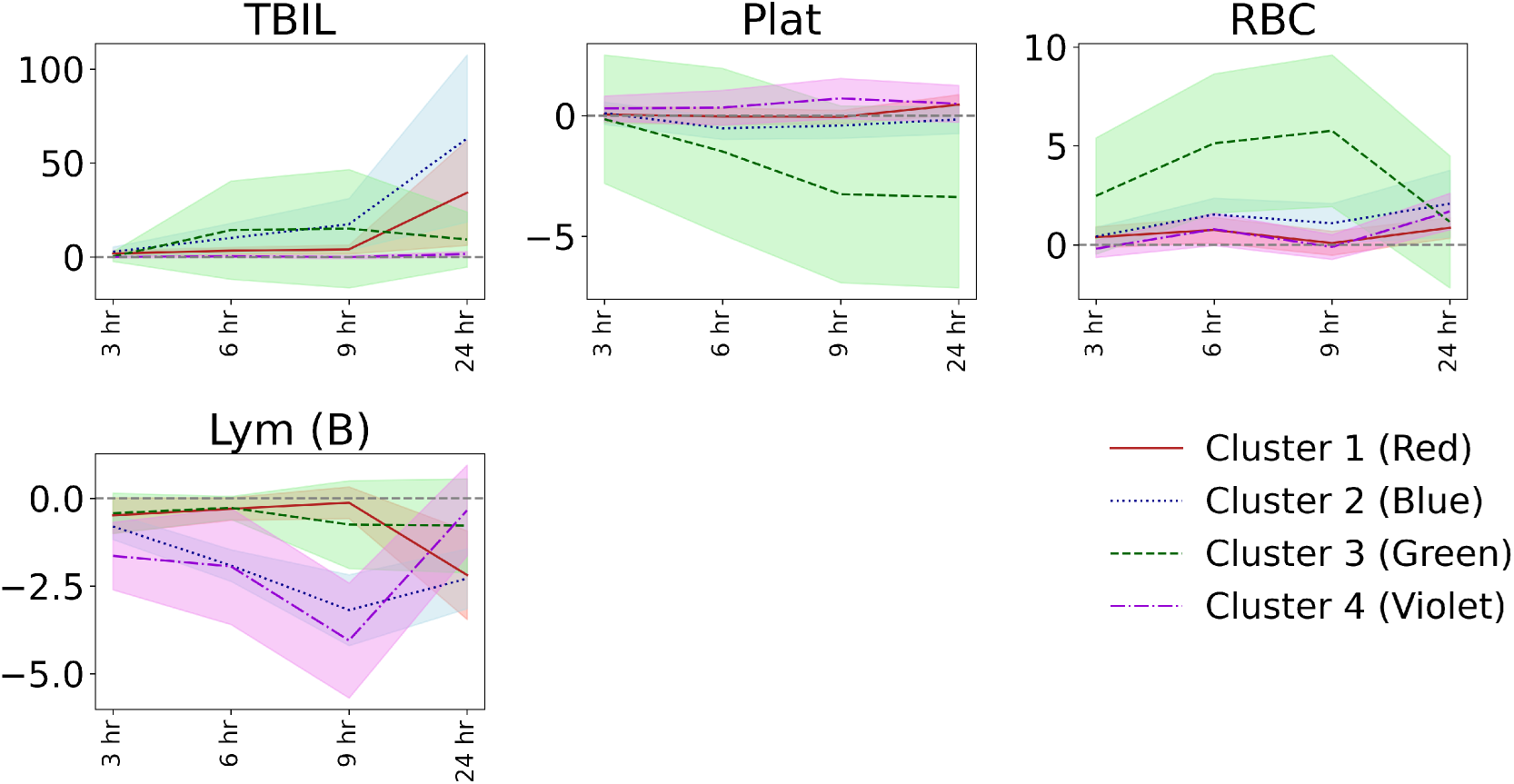
Time course of changes in blood biochemistry. The line is the average value for each cluster, and the filled color around it indicates the 95% confidence interval. Each color corresponds to the color of the compound stratified by analysis of immune cell trafficking. Note that the scale of the x-axis does not correspond to the actual times. TBIL, total bilirubin; Plat, Platelets; RBC, red blood cells; LymB, lymphocytes in the blood.

**Supplementary Figure S12.**
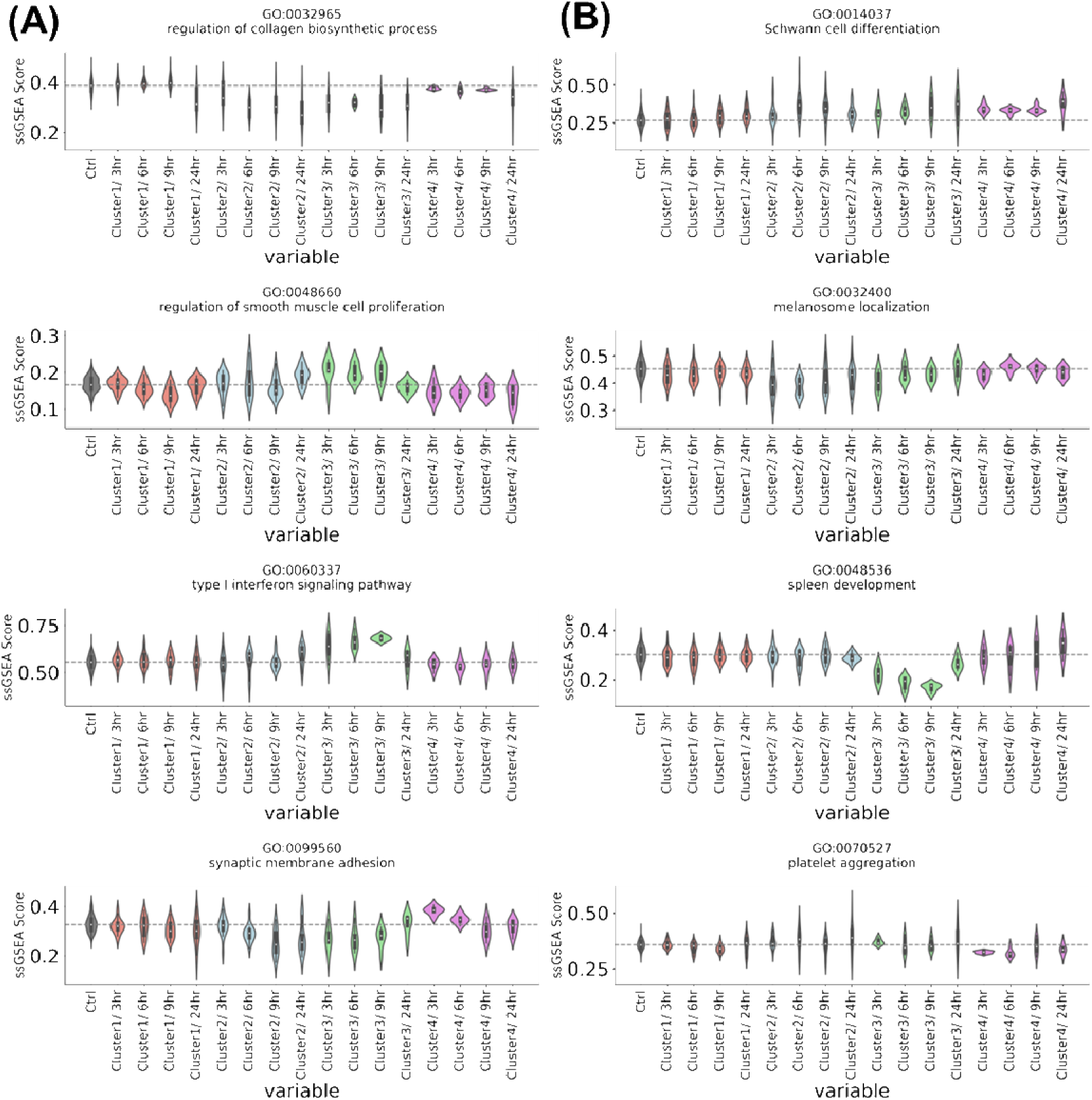
Time course of changes in single sample gene set enrichment analysis (ssGSEA) scores of the liver transcriptomes. For each cluster, calculated from the immune cell trafficking, from the top to the bottom, the gene ontology terms with the lowest p-value in the comparison test with the other clusters are shown. Terms with (A) highest and (B) lowest ssGSEA score in that time compared with other clusters.

**Supplementary Figure S13.**
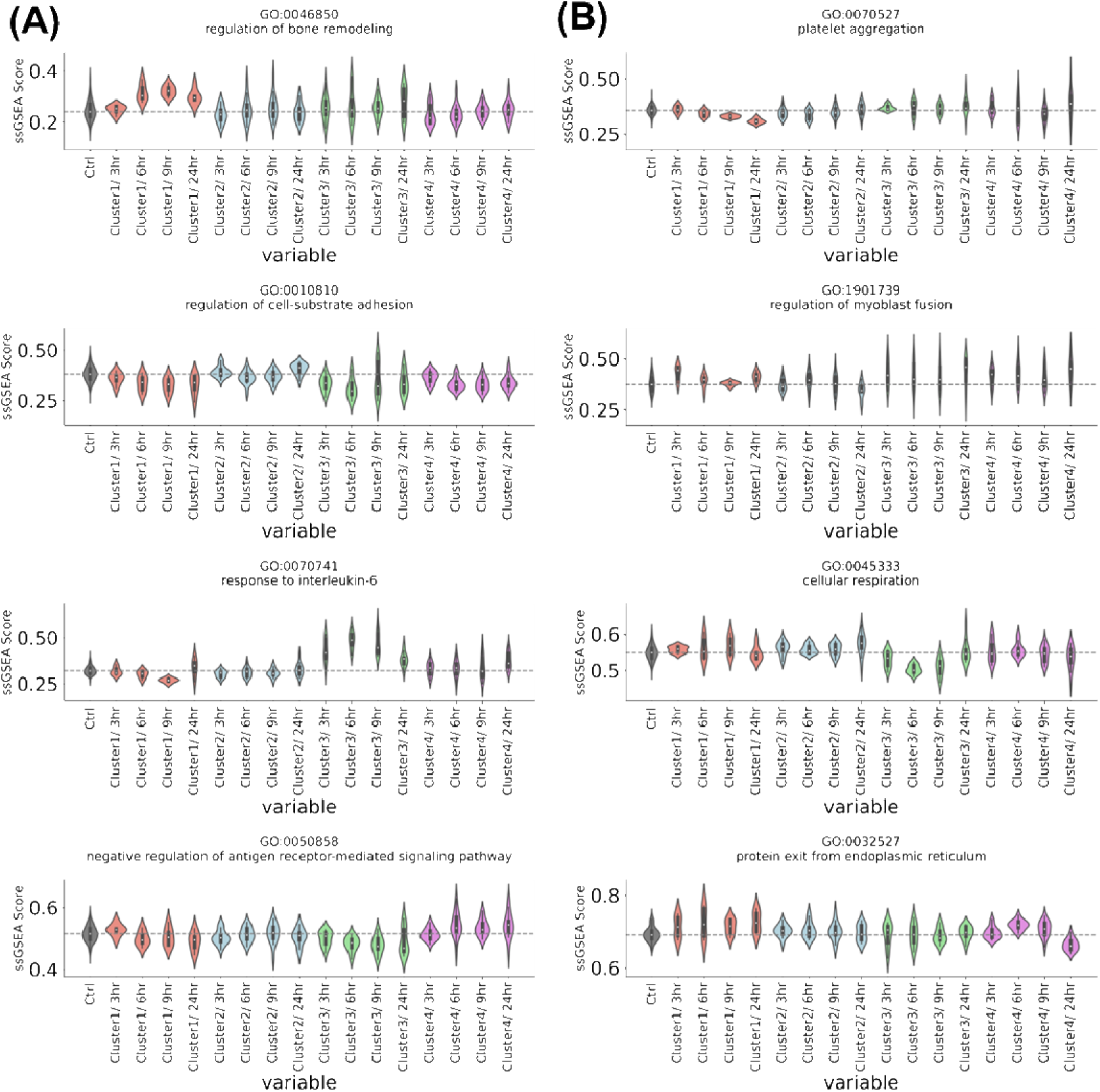
Time course of changes in single sample gene set enrichment analysis (ssGSEA) scores of the liver transcriptomes. For each cluster, calculated from the liver transcriptomes, from the top to the bottom, the gene ontology terms with the lowest p-value in the comparison test with the other clusters are shown. Terms with (A) highest and (B) lowest ssGSEA score in that time compared with other clusters.

**Supplementary Figure S14.**
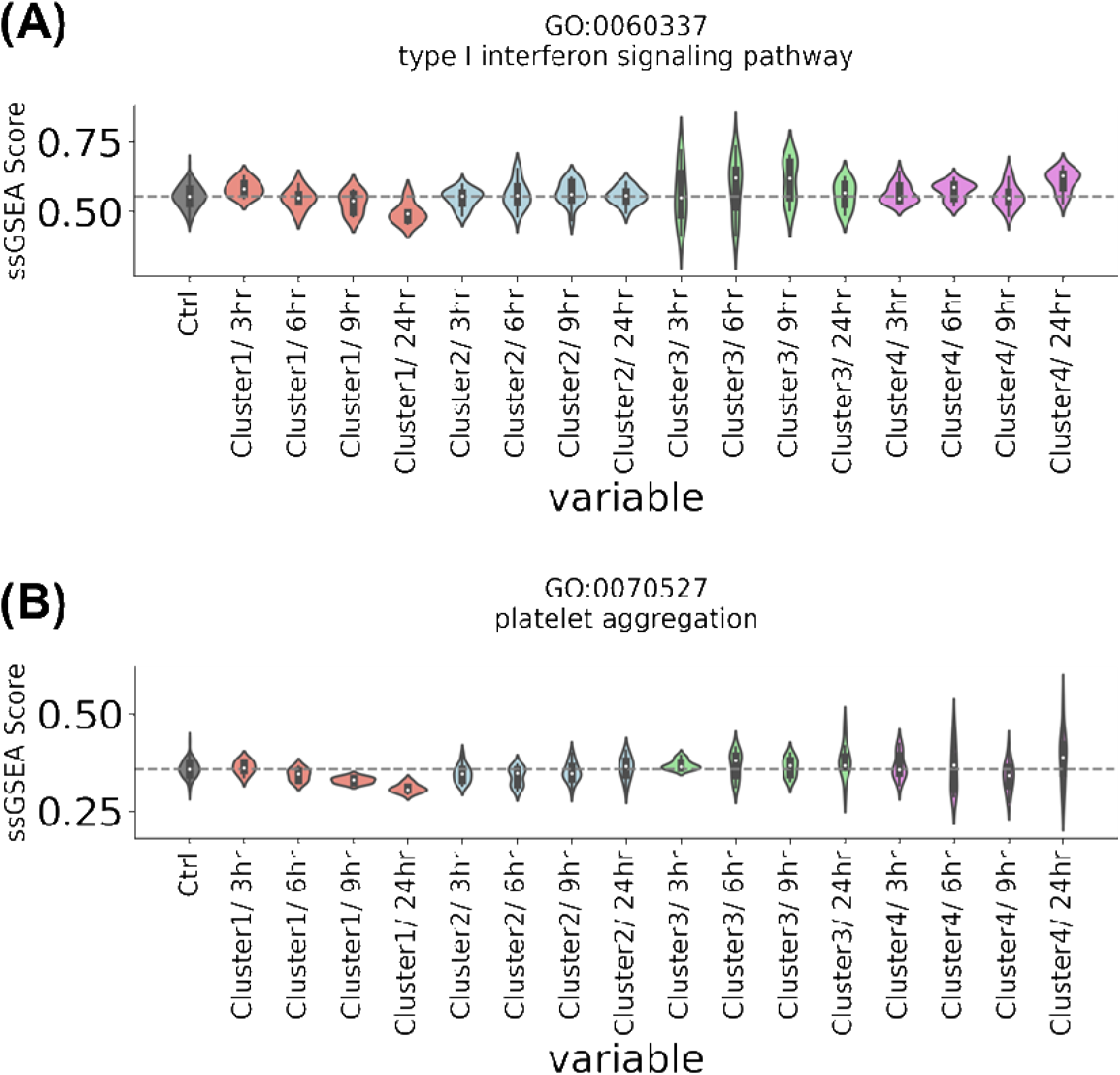
Time course of changes in single sample gene set enrichment analysis (ssGSEA) scores of the liver transcriptomes. For gene ontology terms of (A) interleukin-1-mediated signaling pathway and (B) platelet aggregation.

**Supplementary Figure S15.**
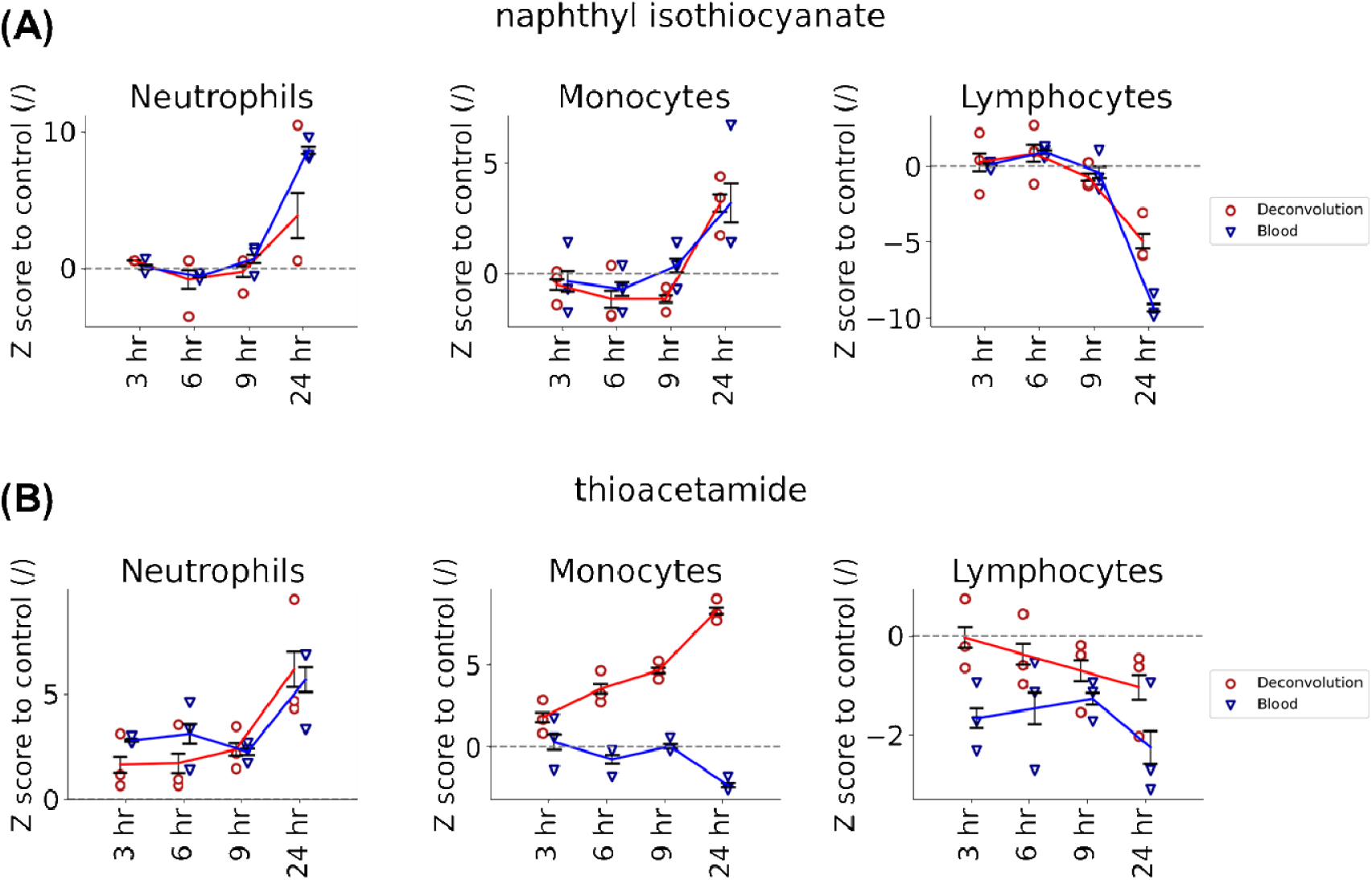
Time course of changes in estimated neutrophils, monocytes, and lymphocytes in the liver using deconvolution and that of measured in the blood of rats administered (A) naphthyl isothiocyanate or (B) thioacetamide as determined from the Open TG-GATEs database. Note that all values were converted to *z*-score for the corresponding control samples. Estimated values for lymphocytes are estimated from sum of NK, B, CD4 T, and CD8 T cells.

